# Deciphering the tumor-specific immunopeptidome *in vivo* with genetically engineered mouse models

**DOI:** 10.1101/2021.06.30.450516

**Authors:** Alex M. Jaeger, Lauren E. Stopfer, Emma A. Sanders, Demi A. Sandel, William A. Freed-Pastor, William M. Rideout, Santiago Naranjo, Tim Fessenden, Peter S. Winter, Ryan E. Kohn, Jason Schenkel, Sean-Luc Shanahan, Alex K. Shalek, Stefani Spranger, Forest M. White, Tyler Jacks

## Abstract

Effective immunosurveillance of cancer requires the presentation of peptide antigens on major histocompatibility complex Class I (MHC-I). Recent developments in proteomics have improved the identification of peptides that are naturally presented by MHC-I, collectively known as the “immunopeptidome”. Current approaches to profile tumor immunopeptidomes have been limited to *in vitro* investigation, which fails to capture the *in vivo* repertoire of MHC-I peptides, or bulk tumor lysates, which are obscured by the lack of tumor-specific MHC-I isolation. To overcome these limitations, we report here the engineering of a Cre recombinase-inducible affinity tag into the endogenous mouse MHC-I gene and targeting of this allele to the Kras^LSL-G12D/+^; p53^fl/fl^ (KP) mouse model (KP; K^b^Strep). This novel approach has allowed us to isolate tumor-specific MHC-I peptides from autochthonous pancreatic ductal adenocarcinoma (PDAC) and lung adenocarcinoma (LUAD) *in vivo*. With this powerful analytical tool, we were able to profile the evolution of the LUAD immunopeptidome through tumor progression and show that *in vivo* MHC-I presentation is shaped by post-translational mechanisms. We also uncovered novel, putative LUAD tumor associated antigens (TAAs). Many peptides that were recurrently presented *in vivo* exhibited very low expression of the cognate mRNA, provoking reconsideration of antigen prediction pipelines that triage peptides according to transcript abundance. Beyond cancer, the K^b^Strep allele is compatible with a broad range of Cre-driver lines to explore antigen presentation *in vivo* in the pursuit of understanding basic immunology, infectious disease, and autoimmunity.

## Main

The success of cancer immunotherapy has led to an explosion of interest in understanding how the immune system recognizes cancer cells^1,2^. Detailed mechanistic investigation in pre-clinical models and patient samples has revealed that responses to immunotherapy are dependent on the presentation of peptide antigens on major histocompatibility complex Class I (MHC-I)^1,3^. MHC-I is a heterotrimeric complex consisting of a heavy chain (H2-K and H2-D in C57BL/6 mice, HLA-A,B,C in humans), a light chain (beta-2-microglobulin, B2m), and a peptide, generally 8-11 amino acids in length. Peptides presented by MHC-I are derived from the proteolysis of numerous intracellular proteins, giving rise to a diverse array of peptide MHC-I complexes, known collectively as the “immunopeptidome”^4^. Expression of mutant proteins, such as those present in cancer or virally infected cells, results in the presentation of foreign peptides or neo-antigens that CD8 T-cells can recognize to drive anti-tumor immune responses^5^.

The complex mechanisms that regulate MHC-I folding and trafficking, peptide processing and loading onto MHC-I, and the biochemical features of MHC-I bound peptides have been elucidated with extensive contribution from numerous labs^2,6–10^. Notably, mass spectrometry-based analyses of the immunopeptidome have deeply characterized endogenously presented peptides in mouse and human cells and tissues, resulting in improved prediction algorithms for peptide-MHC binding and presentation^6,8,11–13^. In conjunction with exome sequencing, these peptide prediction algorithms can be used to identify patient specific neo-antigens, and are increasingly being utilized for personalized immunotherapy^14,7^.

While these advances have undoubtedly improved our ability to rationally design peptide-specific therapeutic modalities, our understanding of the dynamic, context-dependent immunopeptidome is still lacking. Proteomic studies interrogating the immunopeptidome have been carried out almost universally using antibody or affinity immunoprecipitation of MHC-I from cells grown in culture, which lack the microenvironmental and/or tissue-specific stimuli that are sure to impact the repertoire of peptides presented by tumor cells^6,11,12^. Significantly less focus has been paid to understand the immunopeptidome *in vivo*, where existing studies generally profile bulk tumor lysates, which contain contaminating material from tumor infiltrating immune cells and other stromal cells and fail to decipher the cell type(s) that present each of the identified epitopes.

The field currently lacks specialized tools to precisely isolate MHC-I peptides from cells or tissues of interest *in vivo*. While immunocompetent *in vivo* modelling of human tumors remains rudimentary, genetically engineered mouse models (GEMM), including the Kras^LSL-G12D/+^; p53^fl/fl^ (KP) model, represent a tractable system to study the full complement of cellular interactions that drive tumor antigen presentation *in vivo*^15^. GEMMs empower the study of tumor progression in the native tissue microenvironment and recapitulate many of the histopathological features of human cancer. Therefore, GEMMs represent an underappreciated tool to interrogate the tumor immunopeptidome *in vivo* at distinct stages of tumor progression to uncover dynamic features of tumor antigen presentation that have thus far remained elusive^4,16^.

### Design and validation of a Cre-inducible affinity tag onto H2-K^b^ in the KP mouse model

To enable cell type-specific MHC-I purification *in vivo*, we engineered an exon encoding the highly specific affinity tag, StrepTagII, flanked by two pairs of non-compatible Lox sites into intron 1 of the H2-K^b^ locus (Fig. 1a, K^b^Strep). This design results in Cre-dependent inversion of the conditional exon, incorporation of the StrepTag between exon 1 (encoding the H2-K^b^ signal sequence) and exon 2 (encoding the amino-terminus of H2-K^b^), thus enabling Cre-conditional expression of the StrepTag onto the mature H2-K^b^ protein (Fig. 1a). We targeted the K^b^Strep allele to embryonic stem cells derived from mice harboring the Kras^LSL-G12D/+^; p53^fl/fl^ (KP) genotype, in which Cre recombinase activates oncogenic Kras^G12D^ with simultaneous deletion of p53 to initiate autochthonous tumors^15^. Thus, the KP; K^b^Strep mouse enables specific purification of MHC-I complexes from autochthonous tumors *in vivo* (Fig. 1b, Extended Data Fig. 1a-b).

**Figure 1.**
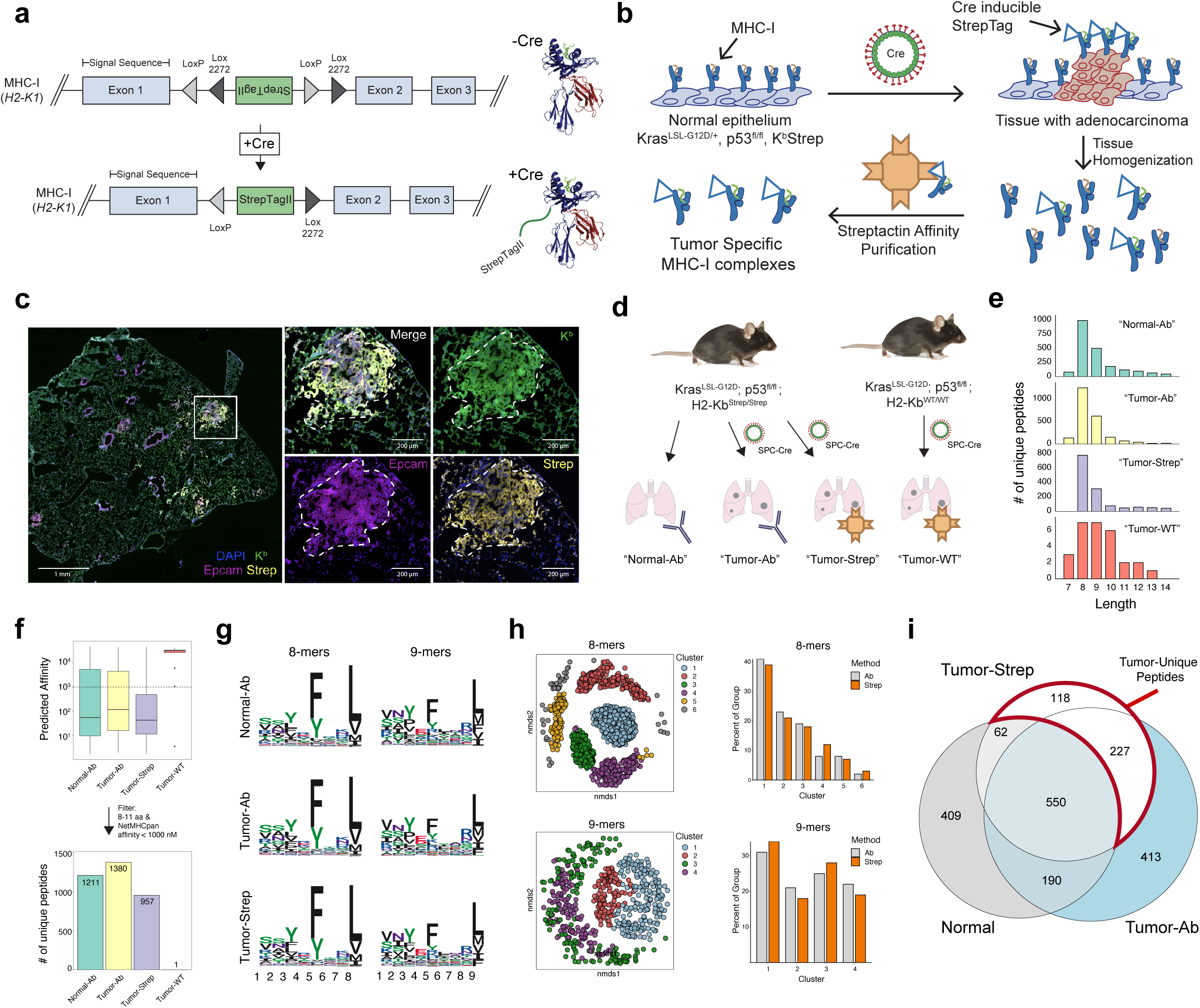
Design and Validation of the KP; K^b^Strep Mouse Model. a) A Cre invertible exon encoding for the StrepTagII epitope was inserted into intron 1 to the H2-K1 gene (left). Cre-recombination induces incorporation of the StrepTag onto the amino terminus of MHC-I (right). b) Schematic illustration depicting how Cre activation of K^b^Strep enables tumor specific isolation of MHC-I in autochthonous tumors. c) Multiplex immunofluorescence of a representative KP; K^b^Strep lung 8 weeks post tumor initiation. White box indicates the zoomed region on the right. d) Experimental schematic depicting the different *in vivo* sample types to be compared in downstream analyses. e) Length distribution of peptides isolated from healthy or KP; K^b^Strep tumor bearing lung with anti-H2-Kb antibody (Normal-Ab, and Tumor-Ab), or KP; K^b^Strep tumor bearing lung with Streptactin affinity purification (Tumor-Strep), or “wild type” KP tumor bearing lung with Streptactin affinity purification (Tumor-WT). f) Number of unique peptides identified in each sample type after filtering for length (8-11 amino acids) and NetMHCPan predicted affinity (<1000 nM). g) Peptide motifs of 8- and 9-mers isolated from Normal-Ab, Tumor-Ab, and Tumor-Strep samples. h) (left) Non-metric multidimensional scaling (nmds) plots depicting clusters of 8- and 9-mer peptides identified from Normal-Ab, Tumor-Ab, and Tumor-Strep samples. (right) Histogram showing the distribution of unique peptides from antibody (Ab) and Streptactin affinity purification (Strep) methods across peptide clusters identified with nmds analysis. i) Venn diagram showing the relationship between peptides identified in healthy lung (Normal), tumor-bearing lung with antibody (Tumor-Ab), and tumor-specific MHC-I purification with Streptactin (Tumor-Strep). Peptides unique to tumors are outlined in red.

To validate this system, we derived pancreatic organoids from KP or KP; K^b^Strep mice and evaluated the inducibility and expression of the StrepTag following Cre-mediated recombination *ex vivo* (Extended Data Fig. 1c)^17^. We confirmed that the K^b^Strep allele was Cre-inducible, presented on the cell surface, and enabled robust purification of intact MHC-I complexes as evidenced by co-purification of B2m (Extended Data Fig. 1d-i). We next sought to validate the KP; K^b^Strep model *in vivo* (Extended Data Fig. 2a). The tumor specificity of K^b^Strep was confirmed with multiplexed immunofluorescence, where PDAC tumors from KP; K^b^Strep mice revealed specific StrepTag detection on tumor nests in the tumor microenvironment (Extended Data Fig. 2a-c). We also noted widespread detection of “wild-type” K^b^ molecules with the standard Y3 antibody, highlighting the importance of tumor-specific MHC-I purification, especially in PDAC, which generally exhibits low tumor cellularity.

For immunopeptidome profiling, we isolated peptides from cells grown in a monolayer (2D), from orthotopically transplanted KP; K^b^Strep organoids (Ortho) or autochthonous tumors initiated through retrograde pancreatic ductal instillation of adenovirus expressing Cre recombinase under control of a Ptf1a (P48) promoter into KP; K^b^Strep mice (Auto) or KP; K^b^WT/WT mice (Auto-WT)^18^. Using an optimized protocol (See Methods, Extended Data Fig 2d), we extracted peptides with the expected characteristics of K^b^ binders in length, predicted affinity, and amino acid content from all samples except for the negative control Auto-WT samples (Extended Data Fig. 2e-i)^9^.

To further probe the performance of our system, we compared peptides isolated from orthotopic and autochthonous PDAC to those identified in normal pancreas tissue from a previous study using traditional antibody-based immunoprecipitation of H2-K^b^. This analysis revealed numerous tumor-specific peptides (Extended Data Fig. 2j). We queried the expression of genes encoding normal pancreas-derived MHC-I peptides (92 genes), orthotopic PDAC peptides (472 genes), and autochthonous PDAC peptides (77 genes) in single cell RNA sequencing (scRNAseq) data from the Tabula Muris. This confirmed that the K^b^Strep system enriched for a immunopeptidome associated with ductal cells, reflecting the expected cellular phenotype of tumor cells in PDAC (Extended Data Fig. 1k-l)^19^. The normal pancreas gene signature exhibited relatively homogenous expression across all cell types in the pancreas, consistent with the notion that antibody-based purification non-specifically samples all cell types in the tissue. Collectively, these results validate that the KP; K^b^Strep system enables high resolution interrogation of cell type-specific immunopeptidomes *in vivo*.

### Identification of Lung Adenocarcinoma (LUAD) associated MHC-I peptides *in vivo*

We next applied the KP; K^b^Strep model to autochthonous lung adenocarcinoma (LUAD) (Fig. 1c-i). Using traditional antibody-based MHC-I immunoprecipitation, we isolated K^b^ peptides from healthy lung (Normal-Ab), 16-week tumor-bearing lung (Tumor-Ab) and compared the peptides to those captured with Streptactin affinity purification from 16 week, KP; K^b^Strep tumors (Tumor-Strep) or KP; K^b^WT tumors (Tumor-WT) (Fig. 1d). Importantly, we observed immunofluorescent detection of the Strep-Tag specifically on tumor cells and with no detection on other cell types within the tumor microenvironment as measured by multiplexed immunofluorescence (Fig. 1c, Extended Data Fig. a-c). Flow cytometric analysis revealed detection of the StrepTag on the surface of cells with an alveolar type 2 surface phenotype (AT2; CD45-EPCAM+MHCII+), which are known to be a major cell of origin for LUAD and consistent with tumor initiating Cre driven by a surfactant protein C (SPC) promoter selective for AT2 cells (Fig. 1d, Extended Data Fig. 3d-g). Notably, the StrepTag was not detected on other CD45-cells in the microenvironment (Extended Data Fig. 3e,f)^20^.

For normal lung samples, we included peptides identified in two independent replicates from healthy lung in this study as well as K^b^ peptides reported previously from healthy lung using the same method^21^. Peptides isolated in the Tumor-Strep samples yielded length distributions, predicted affinities, and amino acid motifs that were indistinguishable from traditional antibody-based methods (Fig. 1e-h). Significantly, peptide profiles from Tumor-WT samples exhibited no enrichment for K^b^ binding characteristics, with only 1 peptide passing size and affinity filters (8-11 amino acids, <1000 nM NetMHCPan 4.1 predicted affinity), compared with 957 unique peptides in the Tumor-Strep samples (Fig. 1f). We evaluated the biochemical features of the identified peptides using non-metric multidimensional scaling (NMDS), which has been previously shown to identify and cluster similar peptide families in human immunopeptidome data^11^. Applying this analytical tool to the 8- and 9-mers in our data set yielded NMDS profiles with 6 and 4 clusters respectively (Fig. 1h). Comparison of peptides derived from StrepTag samples (Strep) or antibody samples (Ab) revealed no significant differences in the distribution of peptides across clusters, confirming that our method does not skew the biochemical properties of peptides that we isolate from *in vivo* tissue (Fig. 1h). Furthermore, we confirmed that the *in vivo* LUAD immunopeptidome that we captured with our method was highly reproducible across replicates (Extended Data Fig. 3j-k).

Comparison of the identity of peptides derived from the Normal-Ab, Tumor-Ab, and Tumor-Strep samples uncovered 345 peptides that were specific to tumors (Fig. 1i, red outline). While 227 of these peptides were also identified with the traditional antibody approach in tumor bearing lungs, 118 were specific to Tumor-Strep samples, suggesting that cell type-specific isolation of MHC-I can provide deeper coverage of the immunopeptidome for cells of interest. 413 peptides were specific to the Tumor-Ab samples, but given the lack of specificity with this method, we could not conclude whether these peptides were derived specifically from neoplastic cancer cells.

### The immunopeptidome reflects cellular identity *in vivo*

We next evaluated whether *in vivo* immunopeptidomes isolated with different methods capture specific cellular identities or cell states within diverse tissue microenvironments. Gene expression signatures were derived from peptides identified by antibody-based isolation of MHC-I from healthy lung (Normal) or tumor-bearing lung (Ab), or with affinity purification of MHC-I specifically from tumor cells (Strep) and applied to scRNAseq data from healthy mouse lung adapted and reanalyzed from the Tabula Muris project (Extended Data Fig. 4a, b)^19^. Within alveolar type 2 (AT2), club/BASC, and basal cells, which represent cell types of origin for LUAD, we observed strong expression of the Strep signature, consistent with the notion that we enriched the immunopeptidome from cancer cells (Fig. 2a). Direct comparison of the Strep and Normal signatures across all cell types revealed a highly significant enrichment for an AT2 phenotype, which was not as pronounced when comparing Ab to Normal. Given that tumors were initiated with a virus expressing Cre from an SPC promoter the strong enrichment for an AT2 phenotype reinforces the specificity of our system. Remarkably, when we compared the Strep and Ab signatures, we observed a robust enrichment for an AT2 phenotype in Strep samples, whereas immune cells were enriched in the Ab sample (Fig. 2a, far right). This suggests that peptides identified by non-specific, antibody-mediated isolation of MHC-I from bulk tumor lysates were obscured by the capture of peptides from tumor-infiltrating immune cells. More generally, these results suggest that cellular identity is broadly reflected in the immunopeptidome.

**Figure 2.**
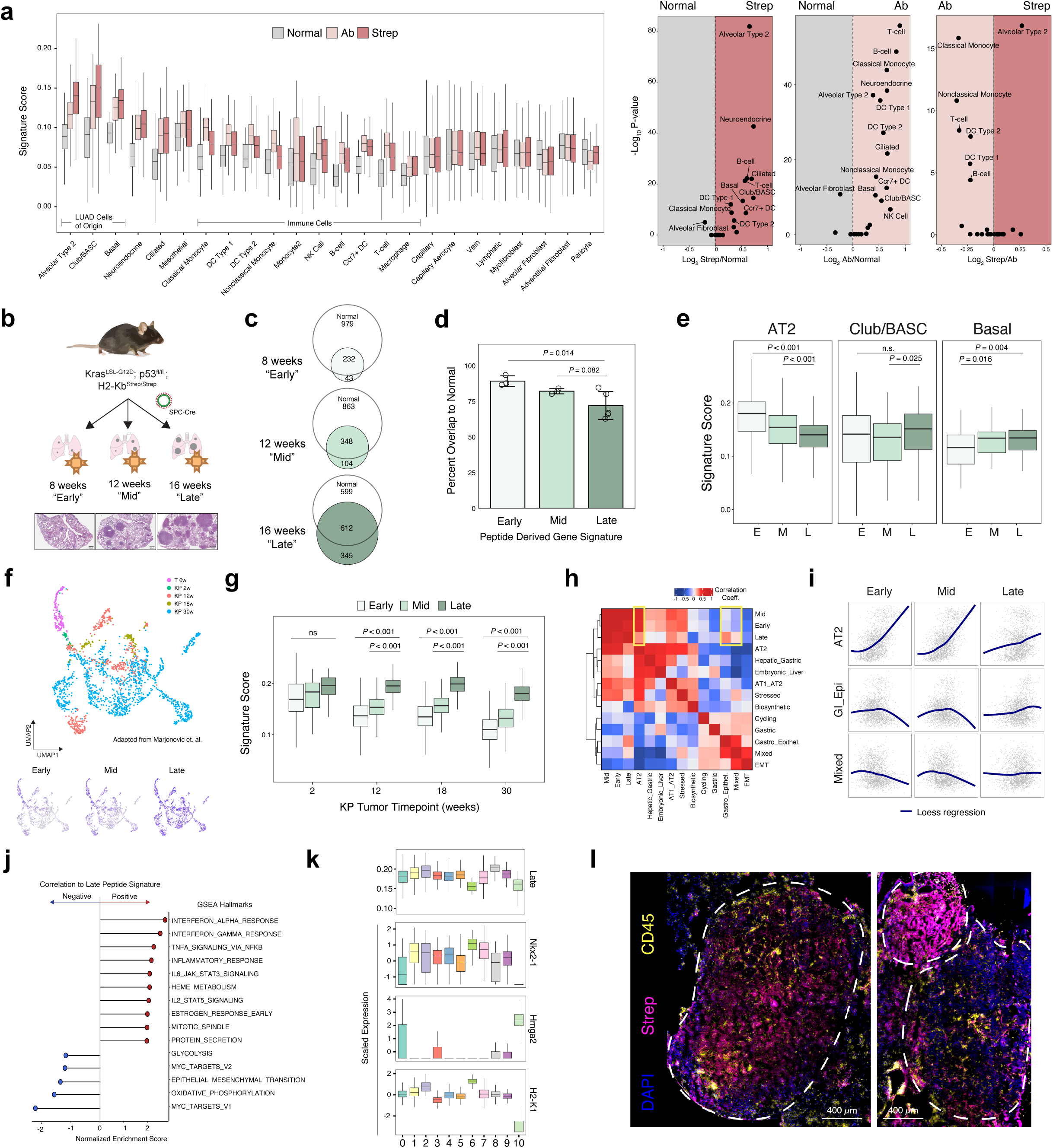
The LUAD immunopeptidome is dynamic and heterogenous throughout tumor evolution. a) Comparison of the relative expression of gene signatures derived from Normal (gray), Tumor-Ab (pink), or Tumor-Strep (red) peptides across all cell types detected by scRNA seq from healthy lung tissue (Adapted from Tabula Muris). Volcano plots for all pairwise comparisons of peptide signatures for each cell type are shown on the right. *P*-value calculated with two-sided Student’s t-test with Bonferroni adjustment. b) Experimental schematic showing the types of samples for comparison of the immunopeptidome through tumor progression. c) Overlap between peptides identified in Early, Mid, and Late stage tumors versus Normal Lung tissue. d) Quantification of the percent overlap from Early (n=3), Mid (n=3), and Late (n=5) stage tumor peptides with normal peptides. Data are mean ± sem. Two-sided Student’s *t*-test. e) Relative expression of gene expression signatures derived from peptides identified in Early (E), Mid (M), and Late (L) stage tumors in alveolar type 2 (AT2), Club/BASC, and Basal cells in the healthy lung. *P* calculated with Mann-Whitney *U* Test. f) UMAP embedding of re-analyzed scRNAseq data from Marjonovic et. al. showing all KP cells throughout tumor progression and expression of the Early, Mid, and Late state peptide signatures (bottom). g) Relative expression of gene expression signatures derived from Early, Mid, and Late peptides throughout KP tumor evolution. *P* calculated with Mann-Whitney *U* Test. h) Correlation of the Early, Mid, and Late signatures versus all gene modules described in Marjonovic et. al. i) Loess regression analysis across all cells scored for the Early, Mid, and Late signatures versus the AT2, GI-Epi, and Mixed modules. j) Gene set enrichment analysis of genes ranked according to their correlation to the Late tumor peptide signature. Gene sets positively correlated to the Late signature are shown in red, and negatively correlated are shown in blue. k) Boxplot showing relative expression of the Late signature, Nxk2-1, Hmga2, or H2-K1 across the 11 distinct clusters found in KP scRNAseq data. l) Multiplexed immunofluorescence depicting tumor specific MHC-I expression (Strep) of a single late stage KP; K^b^Strep tumor (top) or two adjacent tumors (bottom).

While the enrichment of the Strep signature in AT2 cells was encouraging from a methodological perspective, it also prompted us to consider these results in the context of recent work demonstrating the progressive loss of AT2 identity through KP tumor evolution^22,23^. To determine whether the tumor immunopeptidome also evolves to reflect a relative loss in AT2 identity, we evaluated the tumor-specific immunopeptidome at 8 weeks (Early), 12 weeks (Mid), and 16 weeks (Late) following tumor initiation (Fig. 2b, Extended Data Fig. 4c, d). Comparison of MHC-I peptides isolated from Early-, Mid-, and Late-stage tumors to those found in normal lung revealed that throughout tumor evolution the tumor immunopeptidome progressively diverges from normal (Fig. 2c, d). We next derived signatures from genes encoding peptides identified in Early-, Mid-, and Late-stage tumors and applied them to AT2, club/BASC, and basal cell subsets from the healthy lung scRNAseq data. The immunopeptidome signatures exhibit progressive decline in signal within the AT2 compartment, consistent with a loss in cellular identity known to occur during tumor progression in the KP model. In contrast club/BASC cells and basal cells exhibited slightly increased association with Mid- and Late-stage signatures, indicating that as tumor cells lose AT2 identity and adopt alternative cellular phenotypes, a concomitant alteration is observed in the tumor immunopeptidome (Fig. 2e).

### The LUAD immunopeptidome is highly dynamic and heterogeneous through tumor evolution

In an effort to further understand the biological features of peptide profiles in the context of tumor evolution, we applied Early, Mid, and Late peptide-derived gene signatures to published scRNAseq data from normal AT2 cells (T 0w) and KP tumor cells throughout progression (KP 2w, 12w, 18w, and 30w) (Fig. 2f)^22^. We observed progressive enrichment for the Late-stage peptide signature through tumor progression when compared to Early or Mid-stage signatures, uncovering a dynamic nature of the tumor immunopeptidome that has not been previously explored (Fig. 2g). We next queried the correlation of the peptide signatures with previously described expression modules in the KP model^22^ (Fig. 2h). Interestingly, we observed that the Late-stage signature exhibited decreased correlation to the AT2 module and increased correlation to Gastric-Epithelial and Highly Mixed transcriptional programs, previously found to be associated with phenotypic plasticity in late-stage KP tumor evolution (Fig. 2h,i). Moreover, gene set enrichment analysis revealed that gene sets associated with inflammatory cytokine signaling were highly correlated to the Late peptide signature, while gene sets anti-correlated with the Late signature were associated with Myc signaling, metabolic processes, and epithelial to mesenchymal transition (EMT) (Fig. 2j). Overall, this analysis identified extensive heterogeneity in antigen presentation across tumor cell states occurring dynamically in the tumor microenvironment. This was further underscored by examining the metastatic cluster of KP tumor cells in the gene expression dataset, which exhibits very low expression of the MHC heavy chain H2-K1 and was the cluster with the lowest expression of the Late peptide signature (Fig. 2k, cluster 10, low Nkx2-1, high Hmga2, Extended Data Fig. 4e,f)). These observations are also supported by multiplexed immunofluorescence of late-stage tumors revealing significant intra- and inter-tumor heterogeneity in MHC-I presentation that is likely driven by extrinsic (i.e. immune infiltration, nutrient deprivation) as well as intrinsic (i.e. EMT) processes^24^ (Fig. 2l). These results identify extensive heterogeneity in antigen presentation across tumor cell states present in the tumor microenvironment and demonstrate that the immunopeptidome can itself capture the evolution of cell identity throughout LUAD progression.

### Transcriptomic and Proteomic characteristics of Tumor-Unique Peptides

While we noted that gene signatures derived from the totality of the immunopeptidome reflect broad cellular identities as measured by RNA sequencing, we next sought to understand the features of peptides that were *uniquely* presented in KP tumors as compared to healthy lung (Fig. 1i, red outline). Recent evidence suggests that the presentation of MHC-I peptides does not adhere to the law of mass action, and strict concordance between mRNA expression and MHC-I peptide presentation is generally low^4,25,26^. To understand the relationship between transcription and presentation of our tumor-unique peptides, we used published bulk RNAseq data from sorted normal AT2 cells, or cells sorted from Early-, Mid-, or Late-stage KP tumors and examined the expression of genes that encode Tumor-Unique peptides (Fig. 3a)^27^. Genes encoding Tumor-Unique peptides exhibit varied patterns of RNA expression, which did not correlate with either mean mRNA expression or predicted peptide affinity, suggesting that changes in mRNA expression do not exclusively explain tumor-specific presentation of individual peptides. To corroborate this finding, we identified genes that were upregulated at any stage of tumor progression^22^, extracted protein sequences for all gene isoforms, and predicted the affinity of all 8- and 9-mer peptides from this list (Fig. 3b). Ultimately, this resulted in the identification of 17,130 peptides predicted to be enriched in tumor presentation on mRNA expression and predicted affinity alone. However, this pipeline only identified 39 out of 337 total 8- and 9-mers in our Tumor-Unique peptide list, reinforcing the importance of evaluating cell type/tissue specific MHC-I presentation with empirical, mass spectrometry analysis (Fig. 3b).

**Figure 3.**
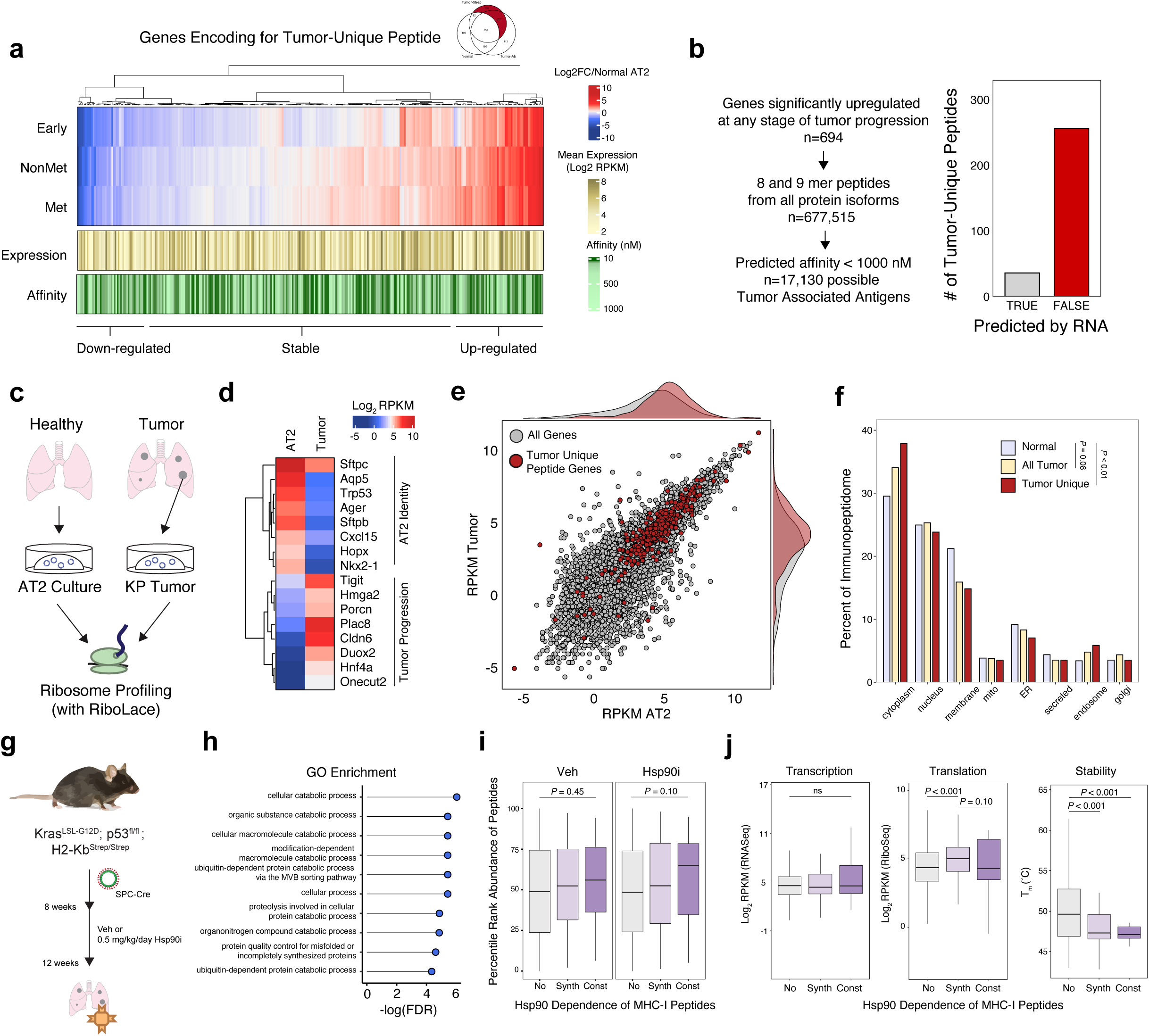
Transcriptomic and Proteomic Features of Tumor-Unique Peptides. a) Heatmap showing the relative RNA expression (red, white, blue), mean RNA expression (gold), and predicted affinity (green) of genes encoding for Tumor-Unique peptides throughout tumor progression compared to normal AT2 cells (adapted from Chuang et. al.). b) Workflow depicting an *in silico* approach to predicting tumor specific peptides based on RNA expression and predicted affinity. Histograms show the number of tumor-unique peptides according to whether they were predicted (grey) or not predicted (red) by RNA/affinity analysis. c) Experiment outline for Ribosome Sequencing by RiboLace. d) Heatmap showing the relative translation intensity (RPKM) for AT2 identity genes and genes associated with KP Tumor Progression. e) Comparison of all genes detected by Ribosome Sequencing based on their relative abundance in normal cells (RPKM AT2) or tumor cells (RPKM Tumor). Genes encoding for tumor unique peptides are highlighted in red. Density plots of the RPKM abundance of all genes (gray) or tumor unique peptide genes (red) are shown for AT2 (top) or tumor (right). f) Subcellular compartment distribution of source proteins for peptides found in Normal tissue (gray), All Tumor peptides (yellow), and Tumor-Unique peptides (red). *P* calculated with Fisher’s exact Test with Monte-Carlo simulation. g) Experimental schematic of KP; K^b^Strep tumor treatment with either vehicle control or 0.5 mg/kg/day NVP-HSP990 prior to tumor specific MHC-I isolation. h) Gene ontology analysis of source proteins for peptides that were only found in Hsp90i treated samples ranked according to FDR enrichment significance. i) Rank ordered abundance of peptides derived from Non-clients (No, gray), synthesis clients (Synth, light purple) or constitutive clients (Const, dark purple) in Veh and Hsp90i treated samples. *P* calculated with the Kologorov-Smirov Test. j) RNA abundance, translation rate, and melting temperatures across non clients (No), synthesis clients (Synth) and constitutive clients (Const). *P* calculated with Mann-Whitney *U* Test.

Recognizing that mRNA expression is mechanistically well removed from the presentation of peptides on MHC-I, we decided to determine whether the relative rate of protein synthesis in tumor or normal cells could better predict the presentation of tumor-specific peptides. To this end, we derived organoid cultures of normal AT2 cells from healthy mice or from a dissected KP tumor and performed ribosome profiling (RiboSeq) (Fig. 3c, Extended Data Fig. 5c, d)^28,29^. Ribosome-protected fragments of mRNA from both normal and tumor samples exhibited characteristics of active translation (Extended Data Fig. 5e, f). Genes associated with AT2 identity (Sftpc, Sftpb, Ager, Aqp5) exhibited higher translation rates in the AT2 samples, while genes associated with KP lung tumor progression (Hmga2, Porcn, Tigit, Hnf4a) were translated at higher rates in tumor samples (Fig. 3d, Extended Data Fig. 5g). However, examination of genes encoding Tumor-Unique peptides revealed similar translation rates in tumor vs normal samples, suggesting that differential translation is also not sufficient to predict tumor-specific MHC-I peptide presentation (Fig. 3e).

The inability of mRNA abundance or translation rate to fully describe the Tumor-Unique immunopeptidome prompted us to consider post-translational features that inform tumor-specific presentation, including the size, stability, and subcellular localization of peptide source proteins^30^. While source protein length and stability were similar across samples, we uncovered a shift towards the sampling of proteins in the cytoplasm and endosome at the expense of sampling the membrane and ER in the tumor immunopeptidome compared to normal tissue (Fig. 3f, Extended Data Fig. 6a, b). This feature of the tumor immunopeptidome was also progressively gained through tumor evolution (Extended Data Fig. 6a). Biophysical characteristics of tumor cells such as surface to volume ratio, or differences in protein degradation pathways, including endoplasmic reticulum-associated degradation (ERAD), may underlie the difference in compartment sampling^31,32^. We also uncovered enrichment of protein families giving rise to the tumor-unique immunopeptidome, including MHC-II antigen presentation (a pathway active in AT2 cells), N-linked glycosylation, oxidative stress, and endosome trafficking (Extended Data Fig. 6c, d). Collectively, these data suggest that rather than transcriptional or translational inputs alone, the tumor immunopeptidome may be influenced by additional post-translational mechanisms.

### Disrupting protein folding pathways alters the tumor immunopeptidome *in vivo*

We next sought to perturb post-translational processes through inhibition of the molecular chaperone heat shock protein 90 kDa (Hsp90), which we previously found to augment MHC-I antigen presentation to stimulate anti-tumor immune responses and alter the tumor immunopeptidome in an interferon independent manner^33^. We treated tumor-bearing KP; K^b^Strep mice with either vehicle or low dose, 0.5 mg/kg/day Hsp90 inhibitor from 8-12 weeks post tumor initiation and isolated the tumor immunopeptidome (Fig. 3g). Hsp90i treatment increased the abundance of functional MHC-I complex that we purified, as measured by B2m co-purification but did not alter the biochemical features of peptides that we isolated (Extended Data Fig. 8a-e). However, we noted an increase in the number of unique peptides identified from known Hsp90 clients in Hsp90i treated samples (Extended Data Fig. 8d). StringDb protein-protein interaction network analysis revealed that Hsp90i-specific peptides were enriched from proteins associated with macromolecular catabolism and protein degradation, consistent with the expectation that Hsp90 inhibition increases protein turnover through the inhibition of protein folding (Fig. 3h)^34^. We next evaluated whether we could observe quantitative differences in the abundance of peptides derived from Hsp90 clients following treatment (Extended Data Fig. 8f,g). Protein dependence on Hsp90 can occur at different stages in their state of maturation^35–37^. Consequently, we found that peptides derived from proteins dependent on Hsp90 after their full maturation (constitutive) were more likely to increase in abundance on MHC-I following Hsp90 treatment when compared to proteins dependent on Hsp90 co-translationally (synthesis) or versus non Hsp90 clients (Fig. 3i)^35^. As a corollary, constitutive clients exhibited the lowest thermal stability of all source proteins identified, suggesting that upon Hsp90 inhibition, thermally unstable, constitutively dependent Hsp90i clients are preferentially shuttled into antigen presentation (Fig. 3j)^38^. Lastly, we compared our Hsp90i peptide dataset against all of our tumor-derived peptides, and any peptide that was found to be presented in any normal mouse tissue^19^. This revealed only 4 peptides that were recurrently presented in Hsp90i samples but not in any other sample queried (Extended Data Fig. 8h, i). Taken together, these data demonstrate that the immunopeptidome is shaped through post-translational mechanisms and, more broadly, underscore the potential of the KP; K^b^Strep model to discover treatment-induced changes in the tumor immunopeptidome *in vivo*.

### Discovery of Novel Tumor Associated Antigens in NSCLC

Tumor-associated antigens (TAAs) result from the differential presentation of peptides on tumor cells when compared to healthy tissue throughout the body. Given the ability to compare the tumor-associated immunopeptidome with the corresponding healthy tissue *in vivo*, we evaluated whether the KP; K^b^Strep model has the potential to uncover novel TAAs^39^. Instead of working “bottom-up” from mRNA to MHC-I peptide, our model enables the “top-down” approach of starting with interesting patterns of *in vivo* peptide presentation followed by examining mRNA expression across all murine tissues to nominate and prioritize TAAs (Fig. 4a, Extended Data Fig. 9a, b). Of 1038 peptides identified at any stage to tumor progression, we identified 121 peptides that were not found to be presented on any healthy tissue in the mouse on either MHC allele (K^b^ or D^b^, Schuster et. al.). Filtering this list for peptides derived from genes that exhibit restricted patterns of expression across healthy tissue led to the prioritization of 15 candidate TAA peptides, compared to a difficult-to-prioritize pool of 17,130 epitopes predicted through mRNA expression alone (Fig. 3b, Fig. 4a) ^2,40^. Surprisingly, we found many peptides that were recurrently presented by tumors *in vivo* (Fig. 4a, purple), despite exhibiting very low levels of mRNA expression. This feature was preserved in human immunopeptidome data, where among the 121 TAA human homologs, 42 genes were expressed < 10 TPM in LUAD TCGA, and 13 were sources for peptides presented on A549 cells *in vitro* (Fig. 4b). For prioritized TAAs, we found potential cancer testis antigens (Cfap206, Ccdc158, Ift74, Pdzd8, Gk2, Tmem144), oncofetal antigens (Dnaaf5, Rsl1d1, Gpn3, Zfp462) and tissue-restricted antigens (Crlf1, Prdm15, Hif3a, Csf2a, Slc26a4) (Fig. 4c). Consistent with the identification of embryonic antigens, many genes that encode Tumor-Unique peptides are expressed higher in embryonic lung tissue compared to adult lung (Extended Data Fig. 9g). We also evaluated the immunopeptidome of 2D cultured KP; K^b^Strep cell lines and found only 6 of the 15 prioritized TAAs in cultured KP; K^b^Strep cell lines; and in A549 cells, peptides derived from 4 of the 15 human homologs were identified. Interestingly, the source proteins for the TAAs found in 2D were highly overlapping between mouse and human, suggesting that *in vitro* culture selects for an immunopeptidome that does not fully reflect the *in vivo* immunopeptidome (Fig. 4c). Discordance between *in vivo* and *in vitro* immunopeptidomes is also reflected in the sampling of subcellular compartments, where *in vitro* cultured cells sample nuclear proteins to a significantly greater extent (Extended Data Fig. 9f). Analysis of the mRNA expression and translation of genes encoding prioritized TAAs unexpectedly revealed that compared to all detected peptides, these genes are transcribed and translated at lower abundance (Fig. 4d). This analysis would imply that many potentially interesting peptides that are triaged in current prediction algorithms due to low expression may be readily presented on MHC-I. Collectively, these data suggest that *in vivo* interrogation of MHC-I peptides using the KP; K^b^Strep mouse model can uncover unique features of the tumor immunopeptidome that would be overlooked with *in silico* approaches or traditional IP methods and offers the potential to broaden the landscape of TAAs (Fig. 4e).

**Figure 4.**
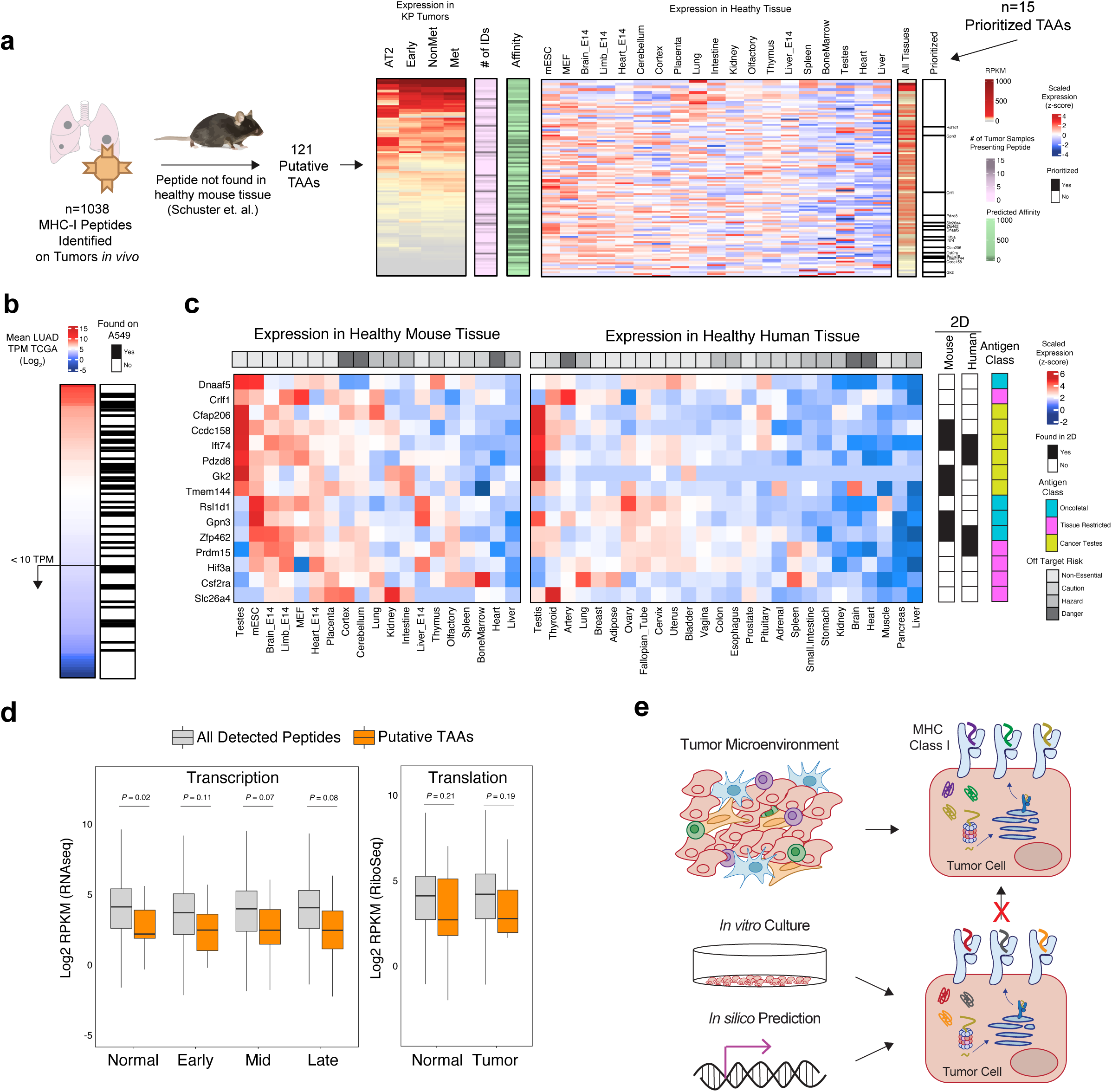
Discovery of novel putative Tumor Associated Antigens (TAAs) in LUAD. a) Workflow for identifying TAAs starting with peptide identification at the mass spectrometry level. 121 peptides were evaluated for tumor expression patterns, recurrent presentation, predicted affinity, and expression in healthy mouse tissue to prioritize 15 TAA candidates. b) Mean RNA abundance of human homologs of the 121 genes in a) across all LUAD samples in TCGA. MHC-I Presentation of a peptide derived from those genes on A549 cells is shown in white/black. c) Scaled RNA expression of genes encoding for 15 prioritized TAAs across all healthy tissues in mouse and human (red, white, blue), whether a peptide was identified from the TAA in the *in vitro* immunopeptidome of KP (mouse) or A549 (human) cells (black/white), tissue Off Target Risk (top, shades of gray), and putative Antigen Class (cyan, pink, yellow). d) Comparison of the transcriptional (RNAseq) and translational (RiboSeq) intensity of genes encoding for all tumor detected MHC-I peptides (grey) or prioritized TAA peptides (orange). 7/15 TAA genes were detected in RiboSeq data with the remaining 8/15 genes having no detectable reads. *P* calculated with Mann-Whitney *U* Test. e) Model depicting the incongruence between the immunopeptidome derived from *in silico* prediction methods, *in vitro* mass spectrometry, and tumor specific *in vivo* immunopeptidomics.

## Discussion

In an effort to expand the toolbox for immunopeptidome-related studies, we developed a novel mouse model that enables the specific isolation of MHC-I complexes from cells of interest *in vivo*. Application of this system to LUAD revealed that the immunopeptidome is highly dynamic through tumor progression and evolves with cellular states adopted by tumor cells. This raises the intriguing possibility that distinct stages of tumor development are associated with an ever-changing landscape of antigen specific T-cells that shift with the dynamic changes in the immunopeptidome through tumor progression.

Interestingly, our results suggest that neither mRNA expression nor translation rate fully explain tumor-specific antigen presentation *in vivo*. While many neo-antigen prediction pipelines include mRNA abundance as a method to triage potential neoantigens, our data suggest that many peptides are recurrently presented *in vivo* despite low transcript levels, highlighting the importance of empirical mass spectrometry evidence to evaluate MHC-I presentation. Further study is needed to elucidate post-translational mechanisms that shape tumor-specific antigen presentation *in vivo*, which may ultimately improve neo-antigen prediction.

TAAs are known to elicit antigen-specific immune responses against tumors^40,41,42^. We found differential presentation of TAAs on tumor cells versus the corresponding normal tissue *in vivo* that could be leveraged with avidity maturation of low affinity T-cell clones that escape negative selection in the thymus. In the context of recent work from our group and others describing insufficient T-cell priming in tumors, it is likely that endogenous responses to these TAAs are limited and may require pharmacological methods to improve priming (e.g. CD40 agonism) and/or vaccination approaches to facilitate CD8 T-cell effector differentiation (Westcott et. al. accepted, Freed-Pastor et. al. accepted)^43^. In the absence of productive endogenous responses, the differential presentation of TAAs identified with this methodology could be leveraged for adoptive cell therapy, antibody drug conjugates, or other therapeutic modalities that require a tumor-specific surface antigen.

We also envision that this model system can be used to evaluate cryptic translation events that result in the presentation of unique MHC-I peptides^29,44,45,46^. Recent studies have found numerous peptides derived from translation outside of canonical coding regions in tumor cells, and, presumably, some of these events may be specific to the *in vivo* context. Additionally, the system reported here will complement the study of post-translationally modified peptides^47^, transposable element-derived peptides^48^, and bacterially-derived peptides^49^ as many of these events will be influenced physiological cues from the *in vivo* microenvironment.

Given the prevalence of cell/tissue specific Cre-driver mouse strains, the K^b^Strep allele presents a novel opportunity to map a high resolution *in vivo* Immunopeptidome Atlas^21^. These data could be cross referenced to other “omic” based tissue atlases to elucidate the relationship between cellular phenotype and the immunopeptidome and to understand how these relationships are disrupted in cancer or other disease states^19^. Additionally, the K^b^Strep allele could be used to evaluate the presentation of viral epitopes during infection, identify auto-antigens in C57BL/6 models of autoimmune diseases, or isolate cross-presented peptides in secondary lymphoid organs.

Our results uncovered aspects of the tumor immunopeptidome *in vivo* that should provoke further investigation into context-specific antigen presentation that will ultimately improve our understanding of tumor-immune interactions. In turn, we describe a versatile tool for the broader research community to interrogate mechanisms of antigen presentation in health and disease.

## Acknowledgements

We would like to thank members of the Jacks Lab for critical comments on the work presented here, especially Zack Ely, Peter Westcott and Nimisha Pattada for conceptual and technical support. We would like to thank George Eng for support with organoid work. In addition, we thank Toni Koller and Richard Schiavoni of the Koch Institute Proteomics core for technical insight on proteomic workflows. Data accessibility from the Tabula Muris was essential for multiple analyses in this manuscript. This work was supported by the Damon Runyon Cancer Research Foundation (A.M.J), an NCI K99 Pathway to Independence Award (A.M.J.), the Howard Hughes Medical Institute, the Johnson and Johnson Lung Cancer Initiative, NCI Cancer Center Support Grant P30-CA1405, the Lustgarten Foundation Pancreatic Cancer Research Laboratory at MIT, the Stand Up To Cancer-Lustgarten Foundation Pancreatic Cancer Interception Translational Cancer Research Grant (Grant Number: SU2C-AACR-DT25-17, W.F.P, T.J.). In addition, we thank the Koch Institute Swanson Biotechnology Center for technical support, specifically the Flow Cytometry, Histology, Preclinical Modeling, Imaging and Testing, and Integrative Genomics and Bioinformatics core facilities.

## Author Contributions

A.M.J. and T.J. conceived, designed, and directed the study. L.E.S. and F.M.W. directed all mass spectrometry analyses. A.M.J., L.E.S., E.S., D.A.S., and R.E.K. performed all experiments. W.F.P. conducted all orthotopic and autochthonous pancreatic surgeries. S.N. provided guidance and reagents for AT2 organoid culture. W. M. R. conducted mESC targeting and chimera generation. A.M.J., L.E.S., T.F., and S.S. performed data analysis. P.S.W. and A.K.S. provided guidance for analysis of scRNAseq data. J.S. and S.L.S. provided technical and conceptual support of the study. A.M.J., L.E.S., F.M.W., and T.J. wrote the manuscript with input from all authors.

## Figure Legends

**Extended Data Figure 1 (Related to Figure 1).**
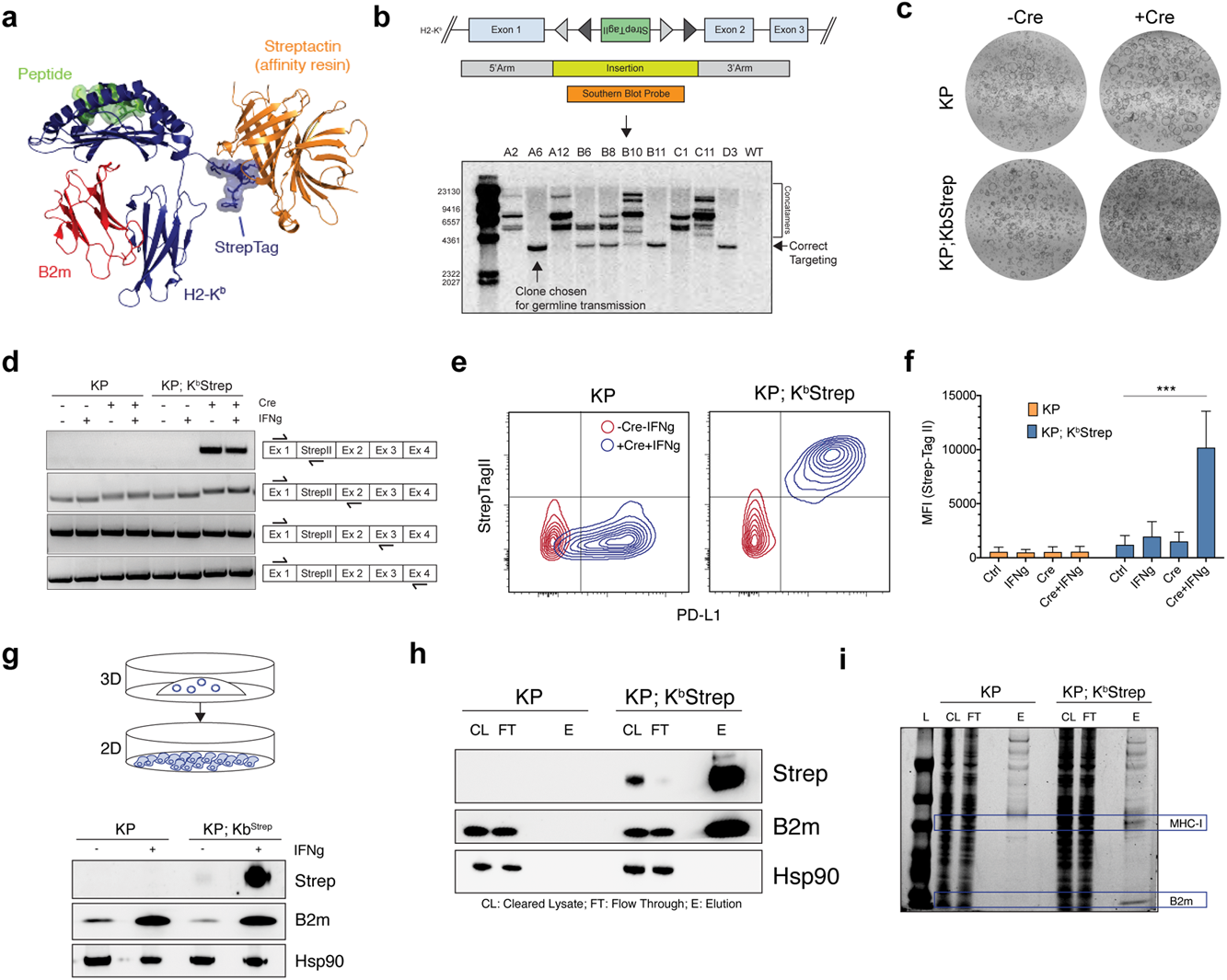
a) Structural model depicting the topology of the K^b^Strep protein during affinity purification. The StrepTag engagement with Streptactin affinity resin does not interfere with peptide or B2m binding. b) Southern blot analysis of K^b^Strep targeted KP* ES cells. c) Brightfield images of KP and KP; K^b^Strep pancreatic organoids pre- and post- Ad-CMV-Cre mediated transformation *ex vivo*. d) RT-PCR analysis of KP or KP; K^b^Strep pancreatic organoids with or without Cre recombination and with or without IFNg treatment. Each row represents a distinct primer set showing no discernable alterations in mRNA splicing with or without StrepTag activation. e) Representative flow cytometry plots detecting cell surface expression of PD-L1 and StrepTagII at baseline (red) and following Cre activation and IFNg treatment (blue). f) Quantification of the median fluorescence intensity of StrepTagII staining in KP (orange) and KP; K^b^Strep (blue) organoids in control, IFNg treated, Cre transformed, and Cre+IFNg treated samples. Data are mean ± sem (n=3). Two-sided Student’s *t*-test. g) Immunoblot analysis of whole cell lysate from KP or KP; K^b^Strep PDAC cells after adaptation to 2D following treatment with IFNg. h) Immunoblot depicting affinity purification of intact MHC-I with Streptactin resin as evidenced by the co-precipitation of B2m. i) Coomassie staining of samples taken from KP or KP; K^b^Strep lysates at various stages of purification. In this experiment, the elution was taken by incubating washed Streptactin resin with SDS-PAGE loading buffer.

**Extended Data Figure 2 (related to Figure 1).**
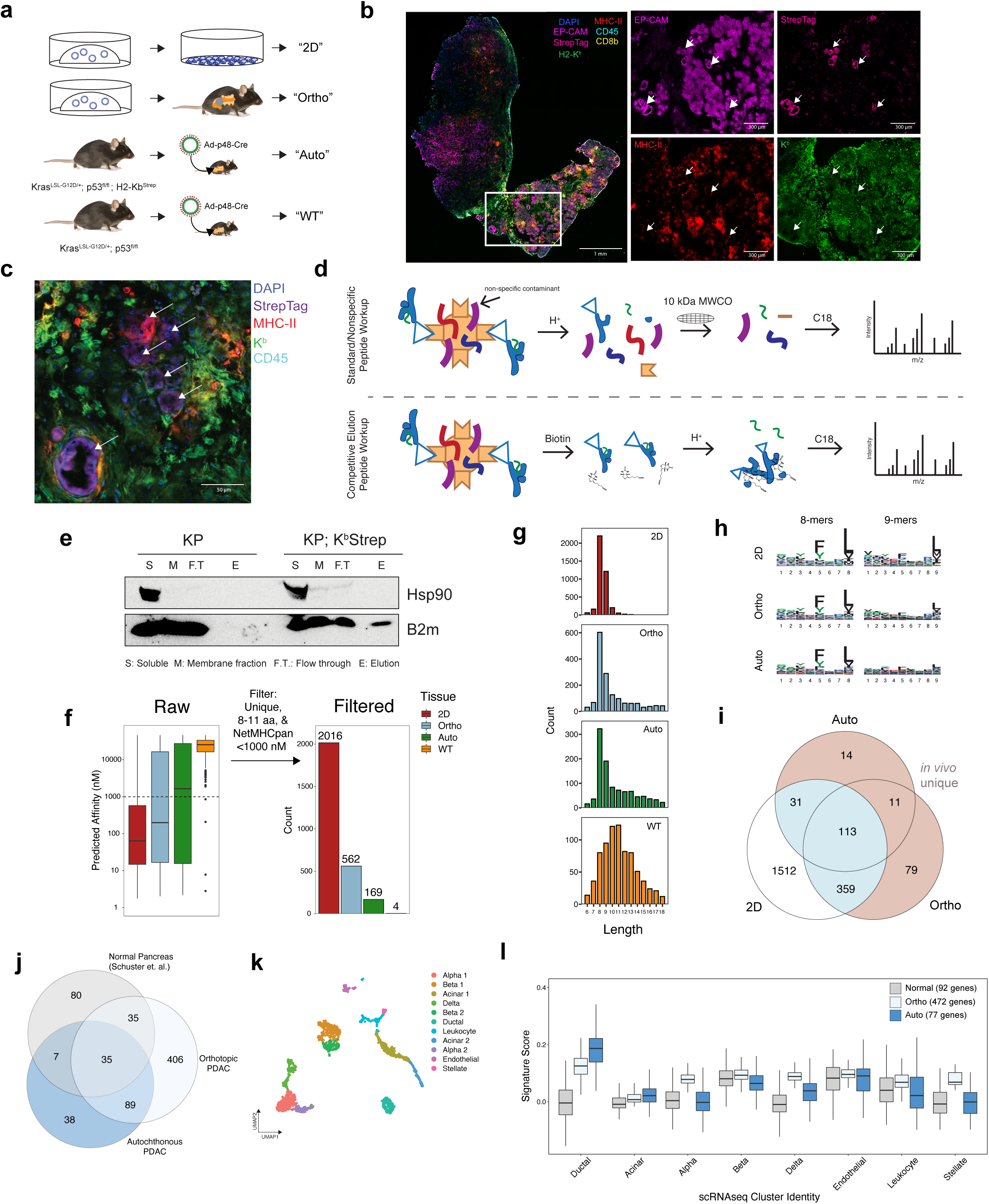
a) Experimental illustration of samples used for immunopeptidome comparison in PDAC. b) Multiplexed immunofluorescence of a representative autochthonous PDAC tumor. White arrows indicate cancer cell nests. c) High magnification multiplexed immunofluorescence image depicting the specificity of StrepTag staining on cancer cells. d) Schematic illustration of the traditional method of peptide extraction (top) and the method used for competitive elution of MHC-I complexes with biotin used in this study (bottom). e) Representative immunoblot demonstrating purification of MHC-I specifically from KP; K^b^Strep tumor bearing mice. f) Number of unique peptides identified in each sample type after filtering for length (8-11 amino acids) and NetMHCPan predicted affinity (<1000 nM). g) Amino acid length distribution of all peptides identified from 2D, Ortho, Auto, or WT samples. h) Peptide motifs of 8- and 9-mers isolated from 2D, orthotopic, and autochthonous tumors. i) Venn diagram comparison of peptides found in 2D, Ortho, or Auto samples. Peptides only found *in vivo* are depicted in tan. j) Venn diagram comparing MHC-I peptides derived from normal pancreas in Schuster et. al. (gray) versus orthotopic transplant (light blue) or autochthonous PDAC (dark blue) in this study. k) UMAP embedding of reanalyzed pancreatic scRNAseq data from the Tabula Muris. For clarity of presentation in Extended Data Figure 1l, cells from a specific lineage were collapsed into a single cluster if they originally separated into multiple clusters (i.e. Alpha 1 and Alpha 2 à Alpha). l) Expression of gene scores derived from normal pancreas peptides (gray), orthotopic PDAC peptides (light blue), and autochthonous PDAC peptides (dark blue) in all cell types of the normal pancreas as measured by scRNAseq from Tabula Muris.

**Extended Data Figure 3 (related to Figure 1).**
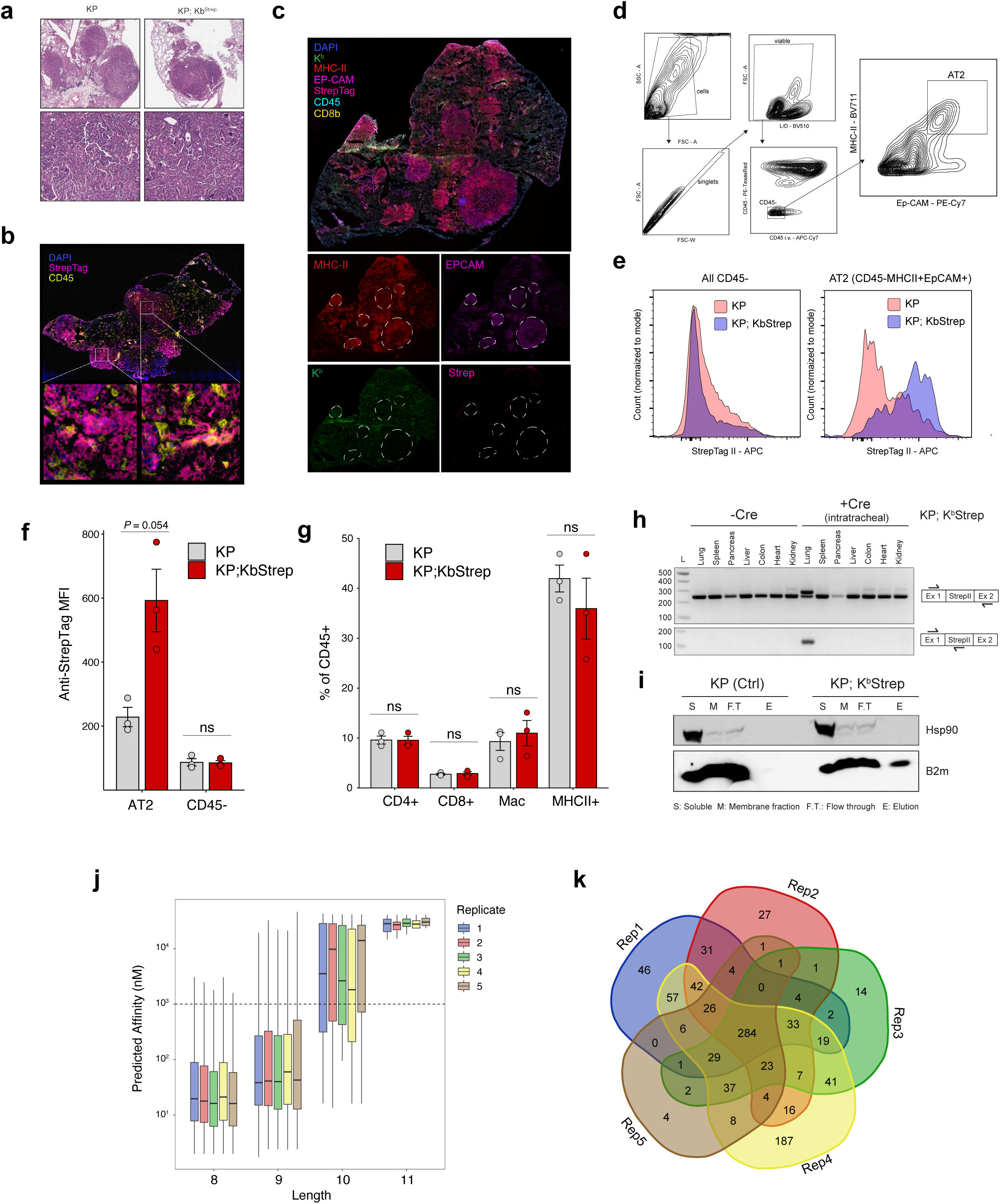
a) Representative H&E images demonstrating adenocarcinoma in KP and KP; K^b^Strep tumors. b) Multiplexed immunofluorescence of late stage KP, K^b^Strep tumors demonstrating specific StrepTag detection on tumor cells within the tumor microenvironment. c) Multiplexed immunofluorescence of a WT KP tumor demonstrating no detection of the StrepTag in tumors outlined in white dotted lines. d) Gating strategy for isolating cells enriched for an alveolar type 2 (AT2) phenotype from KP tumor bearing lung tissue. e) Histograms depicting StrepTagII staining intensity across all CD45-cells (left) or after gating for AT2 cells (right) in KP (pink) or KP; K^b^Strep (purple) tumors. f) Quantification of StrepTag MFI on AT2 or CD45-cells in the tumor microenvironment from KP or KP, K^b^Strep tumors. Data are mean ± sem. Two-sided Student’s *t*-test. g) Relative abundance of CD4 T cells, CD8 T cells, Macrophages, and CD45+MHCII+ immune cells in the tumor microenvironment of KP and KP; K^b^Strep tumors. Data are mean ± sem. Two-sided Student’s *t*-test. h) RT-PCR analysis of diverse tissues in KP; K^b^Strep mice with and without intratracheal Adeno-SPC-Cre administration. Expression of the Strep tagged K^b^ allele is only present in the lung after Cre induction. i) Immunoblot depicting isolation of intact MHC-I complexes as evidenced by co-purification of B2m only from KP; K^b^Strep tissue. j) Distribution of predicted peptide affinity for MHC-I peptides identified in all 5 KP; K^b^Strep replicates. k) 5-way Venn diagram demonstrating the high degree of overlap between 5 independent replicates of extracted MHC-I peptides from 16 week tumors. Each replicate represents 1 tumor bearing mouse.

**Extended Data Figure 4 (Related to Figure 2).**
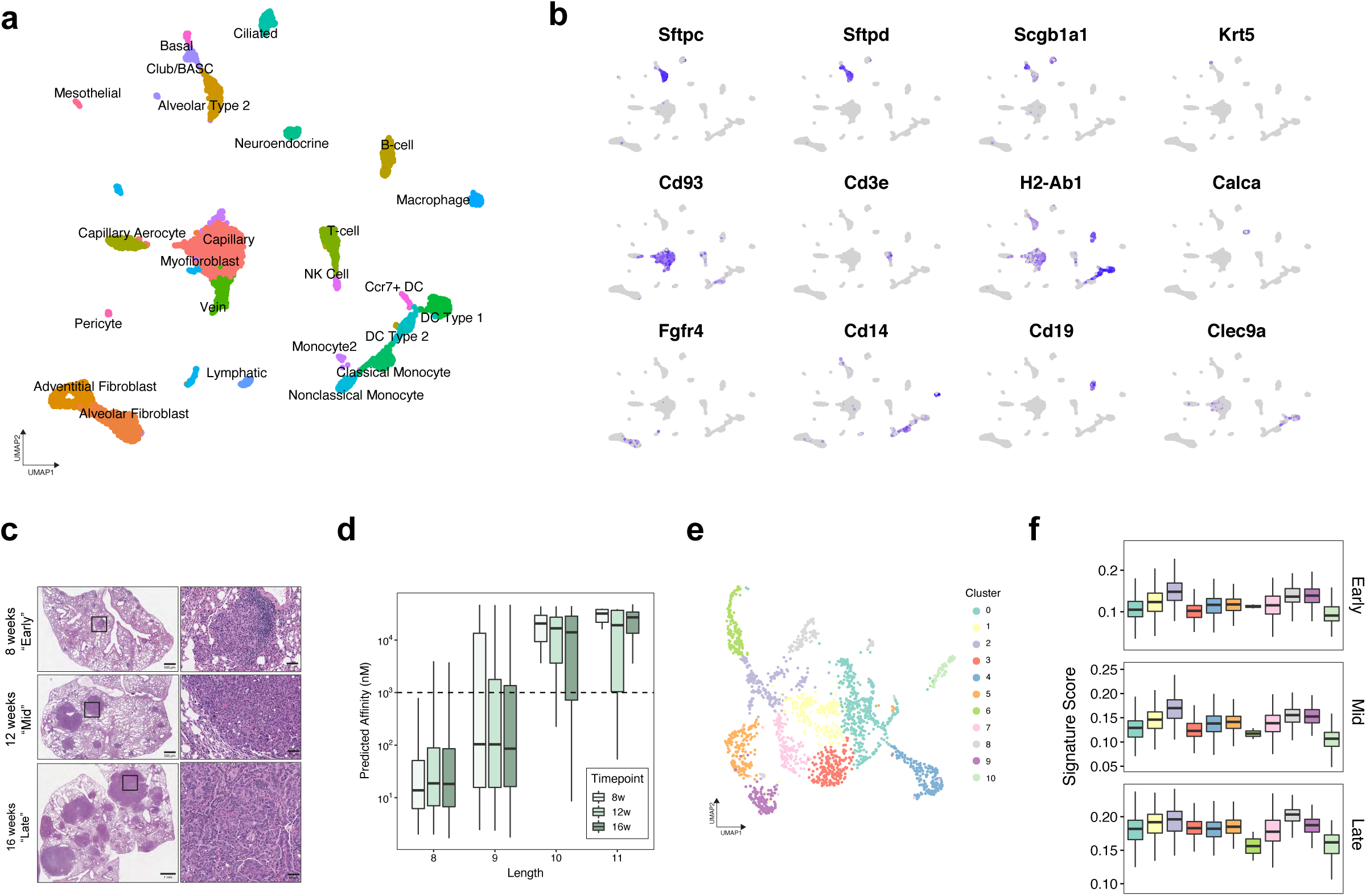
a) UMAP embedding of clusters used for signature expression analysis in Figure 2a. b) Gene expression profiles of cell type marker genes indicating robust clustering of known cell types in the healthy lung. c) Representative H&E stains for Early, Mid, and Late stage tumor samples. d) Length and Affinity characteristics of peptides identified in Early-, Mid-, and Late-stage tumors. e) UMAP depicting clusters used for Figure 2k. f) Expression scores of Early, Mid, and Late gene signatures across all 11 clusters in the KP scRNAseq data.

**Extended Data Figure 5 (Related to Figure 3).**
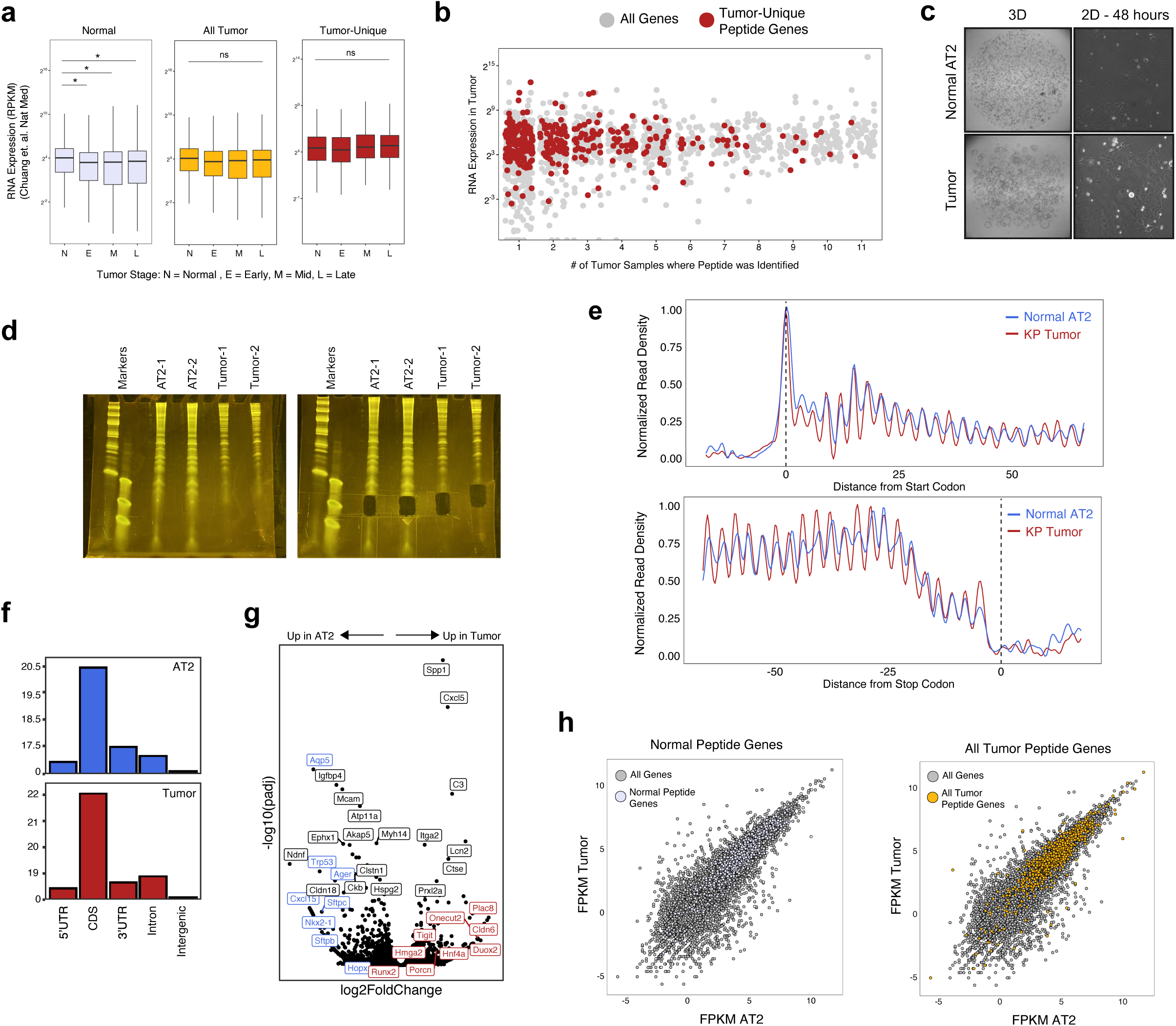
a) RNA Expression of all genes encoding for peptides found in Normal tissue, All Tumor peptides, as well as Tumor-Unique peptides throughout tumor progression (Adapted from Chuang et. al.). *P* calculated with Mann-Whitney *U* Test. b) Relationship between the number of samples that a given peptides was identified (recurrent presentation) versus RNA abundance. c) Representative images of normal AT2 and tumor organoid cultures in 3D organotypic culture or after transient adaptation to 2D monolayer culture prior to RiboSeq processing. d) Denaturing PAGE gels of RNA purified from RiboLace purification. Excised bands used for RiboSeq are indicated on the right and were selected for RNA that was ∼ 30 bp in length. e) Normalized metagene density profiles of reads from normal and tumor cells at translation initiation (left) and termination (right). Both normal and tumor metaprofiles exhibit 3 nucleotide periodicity, indicative of active translation. f) Distribution of uniquely mapped RPF reads across gene features in normal AT2 cells (blue) and tumor cells (red). g) Volcano plot depicting differentially translated genes in tumor versus normal AT2 cells. Data analysis and *P* value calculated with DEseq2. h) Scatter plot of translation rate of all genes (gray), genes encoding for peptides found in normal tissue (purple), or genes encoding for all peptides found in tumor samples (orange).

**Extended Data Figure 6 (Related to Figure 3).**
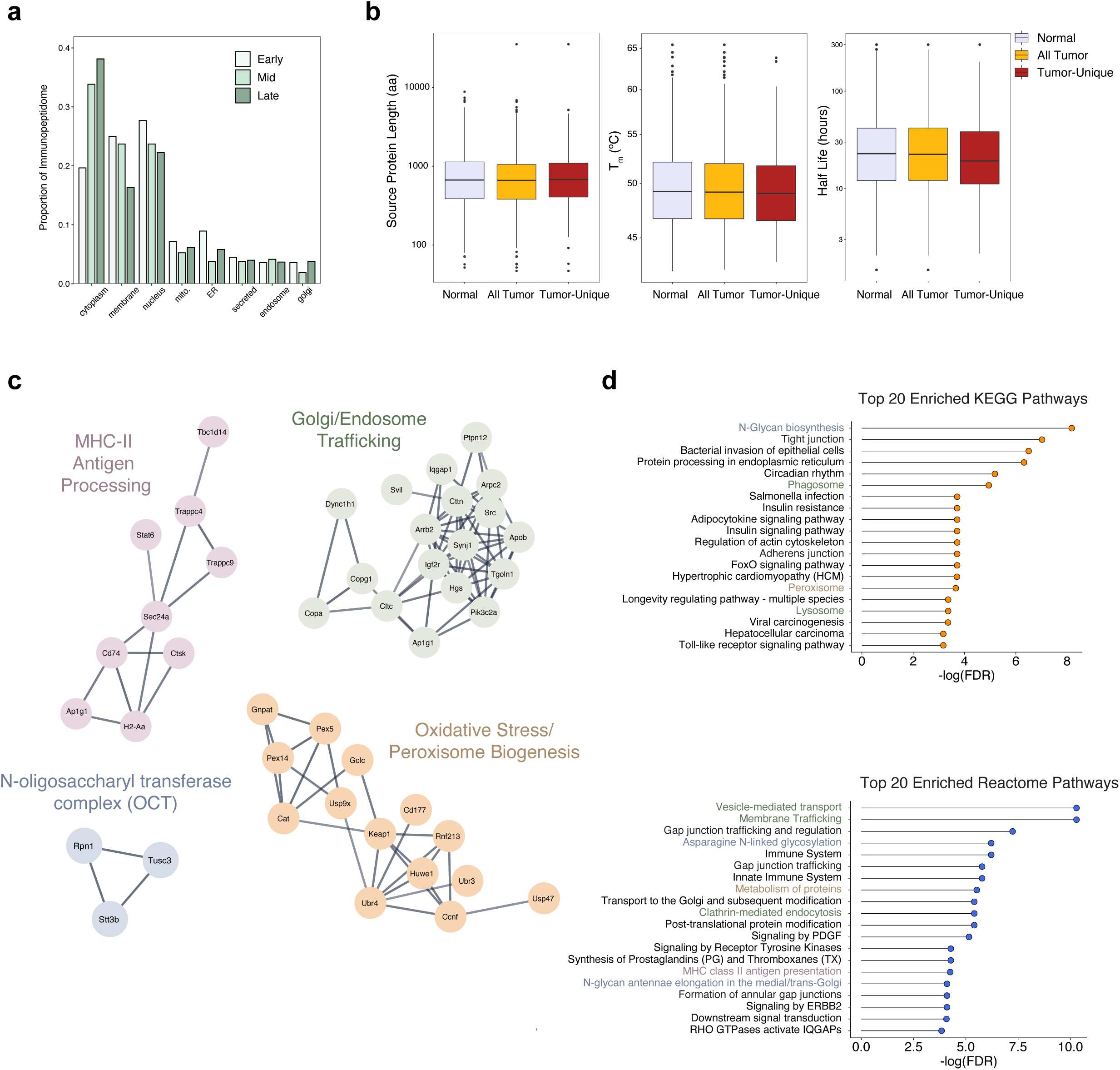
a) Subcellular compartment distribution of source proteins for peptides found in Early-, Mid-, and Late-stage tumors. b) Distribution of Protein length, thermal stability, or protein half-life for source proteins of peptides found in Normal Tissue, All Tumor peptides, or Tumor-Unique peptides. c) StringDb analysis of source proteins for tumor unique peptides indicated in Figure 3i. Clusters of enriched protein families are depicted. d) Gene ontology analysis of tumor unique peptides from KEGG and Reactome databases.

**Extended Data Figure 7 (Related to Figure 3).**
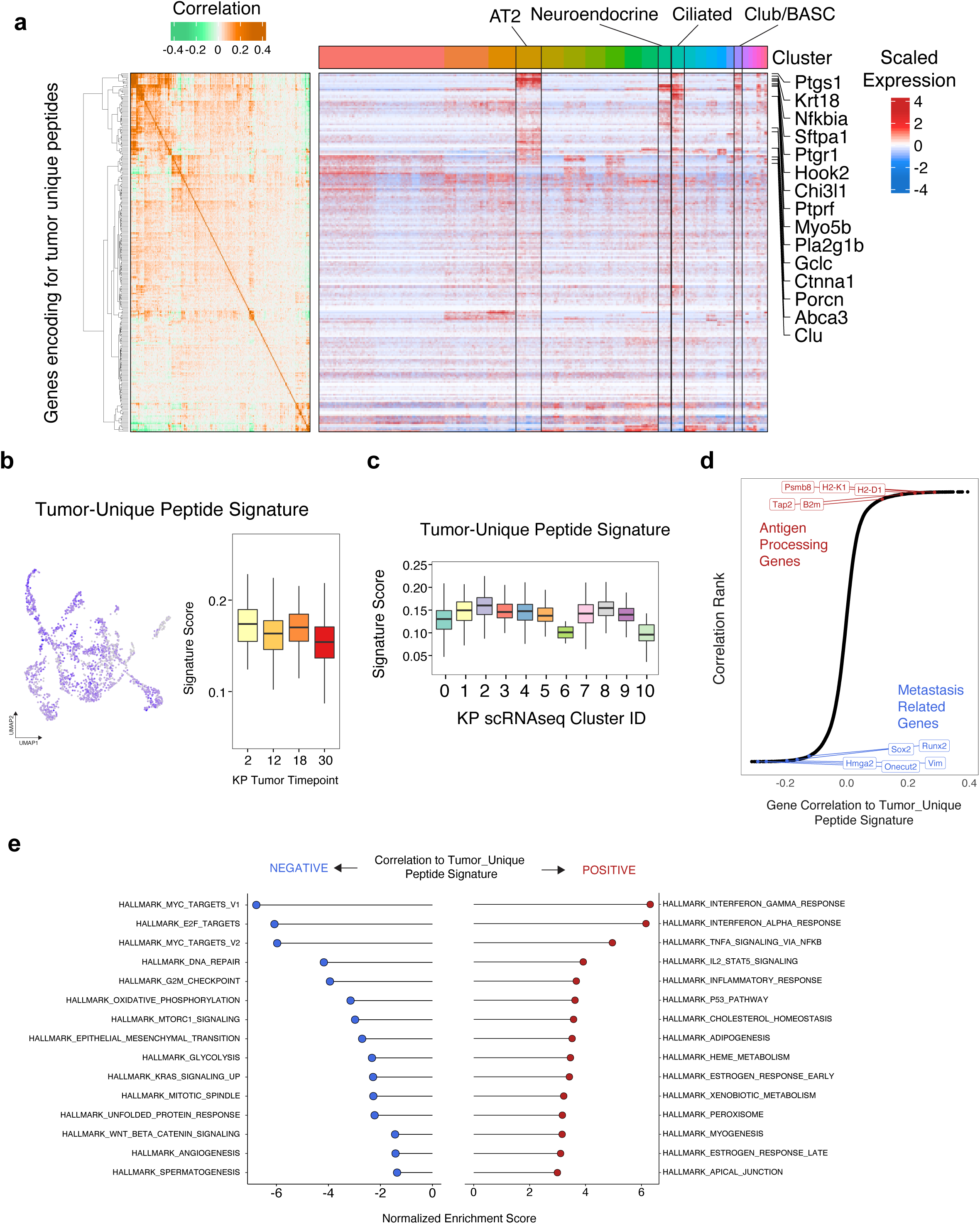
a) Correlogram and corresponding heatmap of genes encoding for Tumor-Unique peptides across all cells in the healthy lung. The majority of genes encoding for tumor-unique peptides exhibit minimal variability between cell clusters except for a select few genes that strongly enrich for AT2, neuroendocrine, Ciliated, and Club/BASC cell clusters. b) Expression of the Tumor-Unique signature across tumor progression in KP scRNAseq. c) Expression of Tumor-Unique peptide signature across clusters in KP scRNAseq. d) Correlation of the Tumor-Unique peptide signature to all genes detected in scRNAseq. Genes related to antigen presentation are highlighted in red and genes related to metastasis are highlighted in blue. e) GSEA analysis of gene sets that are correlated to the Tumor-Unique peptide signature.

**Extended Data Figure 8 (Related to Figure 3).**
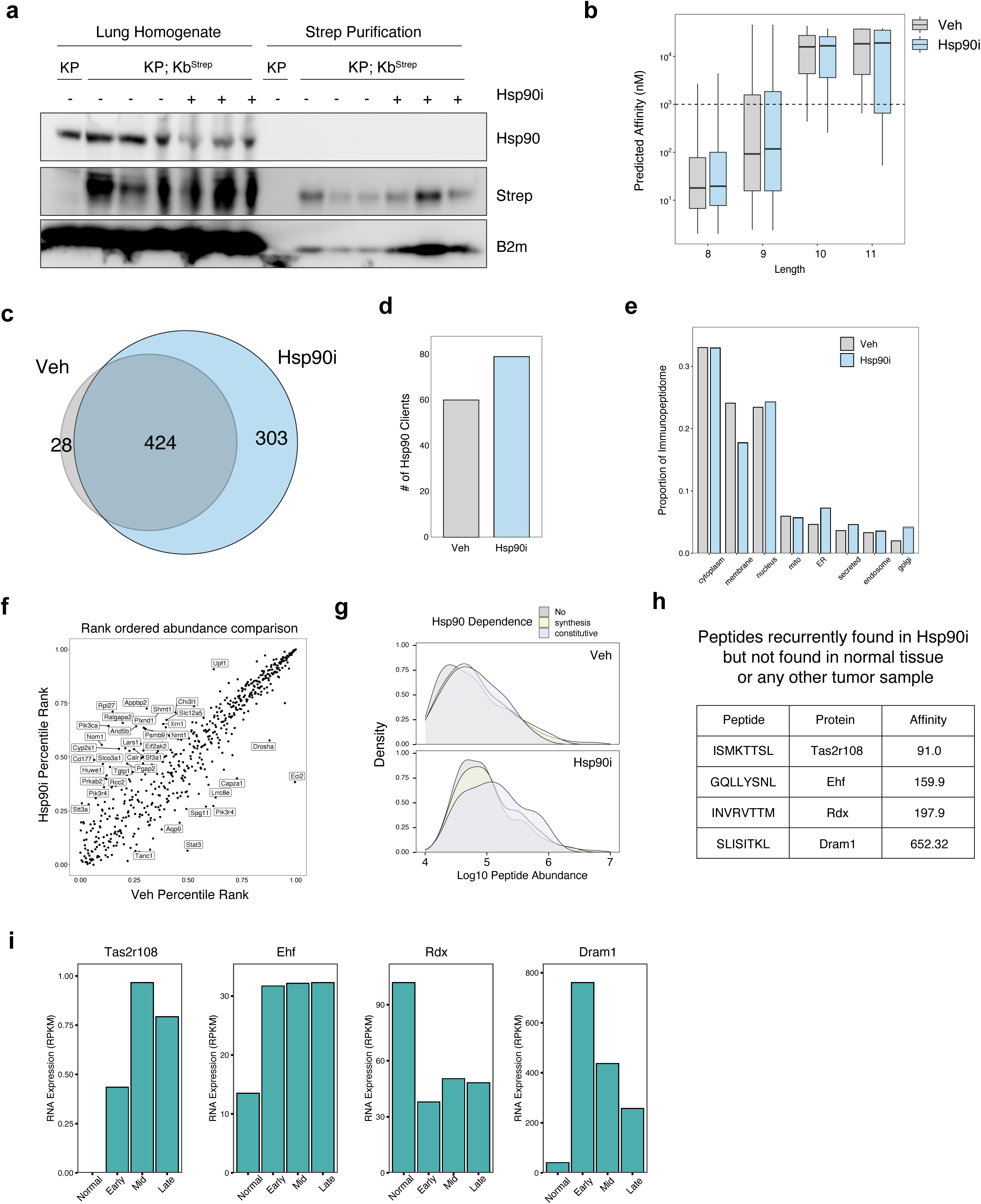
a) Immunoblot analysis of purified MHC-I from Veh and Hsp90i treated tumor samples or KP control tumors. b) Length and affinity distribution of peptides found in Veh (grey) and Hsp90i (blue) treated samples. c) Venn diagram of peptides found in 12 week control tumors or Hsp90i treated samples. d) Number of Hsp90 clients giving rise to peptide in either Veh (grey) or Hsp90i (blue) samples. e) Distribution of peptides identified in Veh or Hsp90i treated samples across subcellular compartments. f) Comparing the rank ordered abundance of all common peptides between Hsp90i treatment and control. g) Density plots of raw peptide abundance for non-clients, synthesis clients or constitutive clients in Veh (top) or Hsp90i (bottom) treated samples. h) Peptides found in Hsp90i treated samples but not in any other tumor sample from any timepoint. i) RNA expression of genes encoding for Hsp90i specific peptides throughout tumor progression as measured by bulk RNA sequencing from Chuang et. al. 2017 Nat. Med.

**Extended Data Figure 9 (Related to Figure 4).**
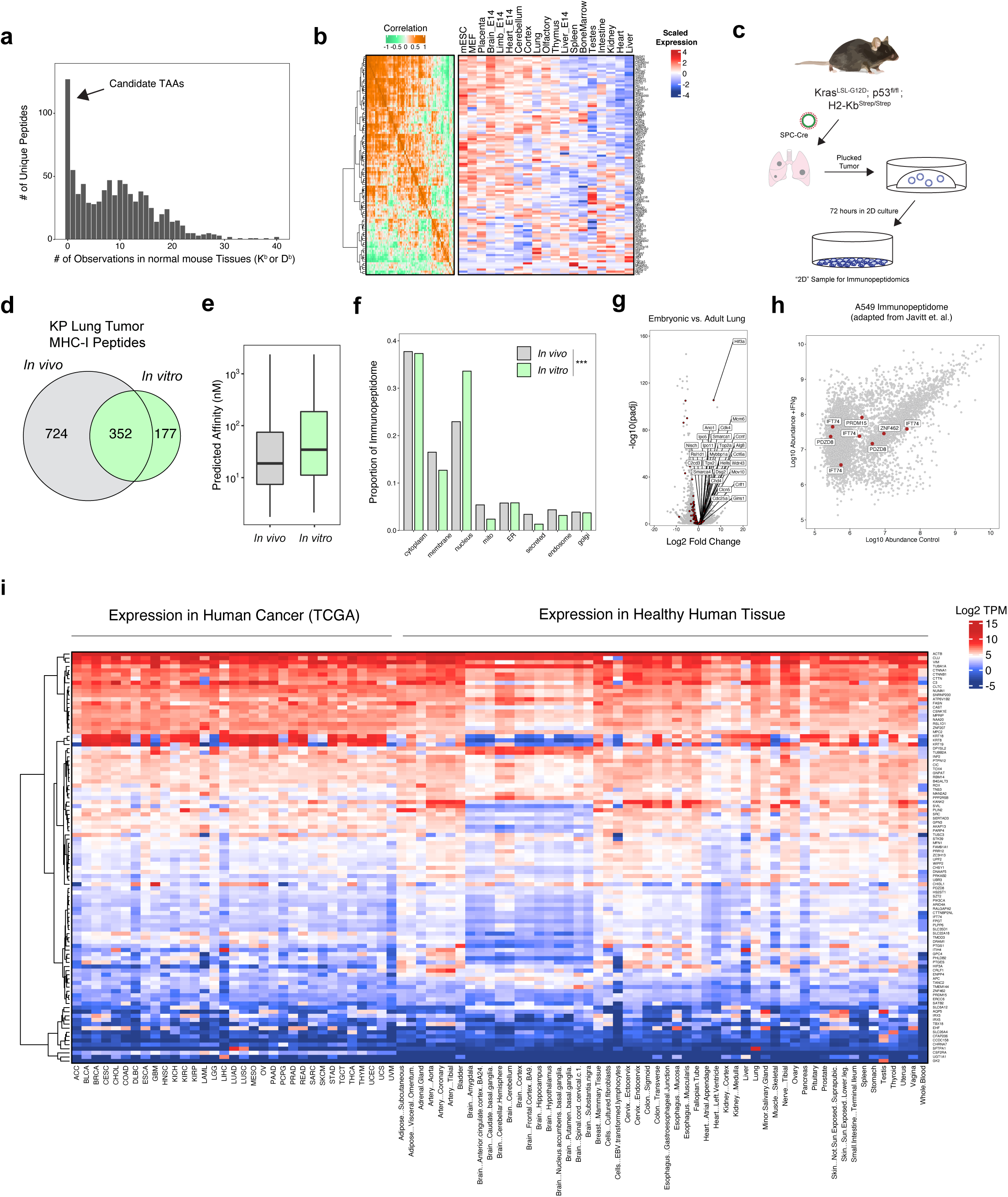
a) Histogram of the number of observations of individual peptides identified in KP tumors across all healthy mouse tissue. b) Correlelogram and heatmap depicting scaled expression of all 121 genes found in Figure 4a. c) Experimental schematic showing the derivation of samples for 2D immunopeptidomics. d) Venn diagram depicting the relationship between peptides identified by KP tumors *in vivo* and those identified *in vitro*. e) Boxplot showing the predicted affinity distributions of peptides isolated *in vivo* and *in vitro.* f) Distribution of source protein subcellular compartments for peptides identified *in vivo* (gray) and *in vitro* (green). *P* calculated with Fisher’s Exact test with Monte Carlo simulation. g) Volcano plot indicating differentially expressed genes between EPCAM+ cells from embryonic day 16.5 and post natal day 28 mouse lung (Adapted from Lung Map Project). Data analyzed and *P* calculated with DEseq2. All genes detected are shown in grey and genes encoding for tumor-unique peptides are shown in red. h) Peptides identified in A549 cells (Javitt et. al.) with and without treatment of IFNg/TNFa. Peptides derived from putative TAA source proteins as identified by KP;K^b^Strep tumors are indicated in red. i) Heatmap depicting the RNA Expression of homologs of potential TAA genes as found in Figure 4a across all individual human tissues and 33 cancer types within TCGA.

## Materials and Methods

### Mice

All animal studies described in this study were approved by the MIT Institutional Animal Care and Use Committee. All animals were maintained on a pure C57BL/6J genetic background, except for ES cell chimeras which were a mix of C57BL/6J and albino C57BL/6J. Generation of Kras^LSL-G12D/+^ and Trp53^flox/flox^ (KP) mice has previously been described (Jackson et al., 2001; Marino et al., 2000).

### mESC culture and CRISPR-assisted gene targeting

Targeted insertion of the Cre-invertible StrepTag allele (K^b^Strep) into the endogenous H2-K1 locus was performed in KP*1 embryonic stem cells, which were generated by crossing a hormone-primed C57BL/6J Trp53^fl/fl^ female with a C57BL/6J Kras^LSL-G12D/+^; Trp53^fl/fl^ male. At 3.5 days post-coitum, blastocysts were flushed from the uterus, isolated, and cultured on a mouse embryonic fibroblast (MEF) feeder layer in ‘ESCM+LIF+2i’ [Knockout DMEM (Gibco), 15% FBS (Hyclone), 1% NEAA (Sigma), 2 mM Glutamine (Gibco), 0.1 mM β-mercaptoethanol (Sigma-Aldrich) 50 IU Penicillin, 50 IU Streptomycin, 1000 U/ml LIF (Amsbio), 3 µM CHIR99021 (AbMole), 1 µM PD0325901(AbMole)]. After 5-7 days in culture the outgrown inner cell mass was isolated, trypsinized and re-plated on a fresh MEF layer. ES cell lines were genotyped for Kras^LSL-G12D/+^;Trp53^fl/fl^, and Zfy (Y-chromosome specific). Primer sequences available upon request. ES cell lines were tested for pluripotency by injection into host blastocysts from albino mice to generate chimeric mice.

DNA mixes containing 3:1 mixes of H2-K1-Strep targeting vector:U6-sgH2-K1-eCas9v1.1-T2A-BlastR were ethanol precipitated prior to Lipofectamine 2000 (Thermo Fisher) transfection of approximately 3 × 10^5^ KP*1 mESCs according to manufacturer’s instructions. Transfected mESCs were plated on MEF feeder cells and selected with 4 ug/mL blasticidin for 48 hours. Cells were then trypsinized and plated at low density onto a fresh plate with MEF feeder cells. After 5-6 days, large colonies were picked using a dissecting microscope and replated into a 96 well plate containing MEF feeder cells. After 4-5 days, each well of the 96 well plate was trypsinized and 1:2 of the material was frozen in ESCM + 10% DMSO and the remainder of the cells were transferred to a 96 well PCR plate for PCR based integration screening. Clones containing homozygous targeting events were then thawed and expanded into 24 well dishes, which were then subjected to genomic DNA extraction and overnight restriction digest with SapI. Digestions were then electrophoresed on a 0.7% agarose gel prior to transfer onto an Amersham Hybond XL nylon membrane (GE Healthcare). Blots were then probed with P^32^-labeled DNA probes comprised of an internal sequence homologous to the genomic insertion containing the StrepTag exon.

Correctly targeted clones were injected into albino C57BL/6 blastocysts. Coat color was used as a surrogate marker for chimerism. Low degree chimeras were chosen for pancreatic organoid generation and high degree chimeras were bred to KP* mice for germline transmission.

### Pancreatic Organoid Isolation

Low degree KP; K^b^Strep chimeras were chosen for organoid isolation using previously described methods (Boj et. al. 2015). Briefly, the pancreas was manually dissected, transferred to a petri dish and thoroughly minced with a razor blade. Minced tissue was then transferred to a 1.5 mL microcentrifuge tube with 1 mL of PBS supplemented with 125 U/mL collagenase IV (Worthington) and DNase (Roche) and incubated rotating at 37 °C for 20-30 min. Cell suspensions were diluted with 9 mL of PBS, and centrifuged at 2000 r.p.m. for 2 min. Cell pellets were then washed with 10 mL of PBS and centrifuged at 2000 r.p.m. for 2 min. The resulting cell pellet was then resuspended in 100% Matrigel (Corning) and plated as 50 µL domes and solidified at 37 °C. Organoids were then cultured in organoid complete media. Purification of cells derived from the targeted ES cells was accomplished with puromycin selection at 6 µg/mL (a puromycin resistance gene is encoded within the LSL cassette upstream of the Kras^G12D^ allele).

Media for pancreatic organoids was formulated based on L-WRN cell conditioned media (L-WRN CM) (VanDussen et al., 2019). Briefly, L-WRN CM was generated by collecting 8 days of supernatant from L-WRN cells, grown in Advanced DMEM/F12 (Gibco) supplemented with 20% fetal bovine serum (Hyclone), 2 mM GlutaMAX, 100 U/mL of penicillin, 100 µg/mL of streptomycin, and 0.25 µg/mL amphotericin. L-WRN CM was diluted 1:1 in Advanced DMEM/F12 (Gibco) and supplemented with additional RSPO-1 conditioned media (10% v/v), generated using Cultrex HA-R-Spondin1-Fc 293T Cells. The following molecules were also added to the growth media: B27 (Gibco), 1 μM N-acetylcysteine (Sigma-Aldrich), 10 μM nicotinamide (Sigma-Aldrich), 50 ng/mL EGF (Novus Biologicals), 500 nM A83-01 (Cayman Chemical), 10 μM SB202190 (Cayman Chemical), and 500 nM PGE2 (Cayman Chemical). Wnt activity of the conditioned media was assessed and normalized between batches via luciferase reporter activity of TCF/LEF activation (Enzo Leading Light Wnt reporter cell line).

Pancreatic organoids were serially passaged with TrypLE Express (Life Technologies). After 4 passages, KP or KP; K^b^Strep organoids were then subjected to *ex vivo* transformation by dissociation, mixing with adenoviral Cre (Ad-CMV-Cre; MOI=500), and re-embedding in Matrigel. After 72 hours, transformants were selected with Nutlin-3a (10 µM, Sigma Aldrich) to select for loss of p53.

### Orthotopic transplantation of pancreatic organoids

Orthotopic transplantation of organoids was performed with minor modifications to previously reported protocols for orthotopic transplantation of pancreatic monolayer cell lines (Kim et al., 2009). Briefly, animals were anesthetized using Isoflurane, the left subcostal region was depilated (using clippers or Nair) and the surgical area was disinfected with alternating Betadine/Isopropyl alcohol. A small (∼2 cm) skin incision was made in the left subcostal area and the spleen was visualized through the peritoneum. A small incision (∼2 cm) was made through the peritoneum overlying the spleen and the spleen and pancreas were exteriorized using ring forceps. A 30-gauge needle was inserted into the pancreatic parenchyma parallel to the main pancreatic artery and 100 μL (containing 1.25*10^5^ organoid cells in 50% PBS + 50% Matrigel) was injected into the pancreatic parenchyma. Successful injection was visualized by formation of a fluid-filled region within the pancreatic parenchyma without leakage. The pancreas/spleen were gently internalized and the peritoneal and skin layers were sutured independently using 5-0 vicryl sutures. All mice received pre-operative analgesia with Bup-SR and were followed post-operatively for any signs of discomfort or distress. Organoid/Matrigel mixes were kept on ice throughout the entirety of the procedure to prevent solidification prior to injection. For orthotopic transplantation, syngeneic C57BL/6J mice (aged 4-12 weeks) were transplanted. Male pancreatic organoids were only transplanted back into male recipients.

### Retrograde pancreatic duct delivery

Retrograde pancreatic duct instillation of lentivirus has been previously described (Chiou et al., 2015). We adapted this technique in a number of ways. Briefly, the ventral abdomen was depilated (using clippers or Nair) 1-2 days prior to surgery. Animals were anesthetized with Isoflurane and the surgical area was disinfected with alternating Betadine/Isopropyl alcohol. A small skin incision was made in the anterior abdomen (∼2-3 cm midline incision extending caudally from the xiphoid process). A subsequent incision was made through the linea alba and incision edges were secured in place with a Colibri retractor. The remainder of the procedure was conducted under a Nikon stereomicroscope. A moistened (with sterile 0.9% saline) sterile cotton swab was used to gently move the left lobe of the liver cranially towards the diaphragm. A second moistened sterile cotton swab was used to gently reposition the colon/small intestine into the right lower abdominal quadrant, until the duodenum was visualized. The duodenum was gently repositioned (still in the abdominal cavity) using moistened cotton swabs until the pancreas, common bile duct and sphincter of Oddi were well visualized. The common bile duct and cystic duct were gently separated from the portal vein and hepatic artery using blunt dissection with Moria forceps. A microclip was placed over the common bile duct (cranial to pancreatic duct branching) to prevent influx of the viral particles into the liver or gallbladder, forcing the viral vector retrograde through the pancreatic duct. To infuse the viral vector, the common bile duct was cannulated with a 30-gauge needle at the level of the sphincter of Oddi and 150 μL of virus was injected over the course of 30 seconds. Gentle pressure was applied at the sphincter of Oddi upon needle exit to prevent leakage into the abdominal cavity. Subsequently, the microclip and Colibri retractor were removed. The peritoneum was closed using running 5-0 Vicryl sutures. The cutis and fascia were closed using simple interrupted 5-0 Vicryl sutures. The entire procedure was conducted on a circulating warm water heating blanket to prevent intra-operative hypothermia. All mice received pre-operative analgesia with sustained-release Buprenorphine (Bup-SR) and were followed post-operatively for any signs of discomfort or distress. For retrograde pancreatic ductal installation, male mice (aged 3-6 weeks) and female mice (aged 3-8 weeks) were transduced with 250,000 TU in serum-free media (Opti-MEM; Gibco).

### Intratracheal Administration of Adenovirus

Adenovirus expressing Cre recombinase from an SPC promoter (Ad5-SPC-Cre, Univ. of Iowa) was prepared by diluting virus stocks to 400 p.f.u./µL into OptiMEM (Gibco) followed by addition of CaCl_2_ to a final concentration of 10 µM. Viral suspensions were then mixed and incubated at room temperature for 20 min before placing on ice. Mice were anesthetized with Isoflurane prior to setting on a custom wire platform to open the mouth. The trachea was then canulated with 22G catheters (Exel) and 50 µL of viral suspension was added to the catheters. Once the mouse aspirated the viral suspension, they were transferred to a prewarmed cage to recover from anesthesia and monitored daily for 3 days to ensure recovery from the procedure. Dilutions of virus were used within 1 hour of preparation.

### Flow cytometry of cultured cells and lung tissue

For staining pancreatic organoids, cells were plated into 4 × 20 uL domes per well in a 12 well plate. Cells were either left untreated or incubated with 20 ng/mL IFNg for 48 hours. At the time of harvest, organoid media was aspirated, and the Matrigel domes were mechanically disrupted with vigorous pipetting in PBS. Cell suspensions were then transferred to 15 mL conical tubes centrifuged at 2000 rpm for 2 min. The resulting cell pellet was then incubated with 1 mL TrypLE Express for 10-15 minutes. The dissociation reaction was quenched with 10 mL PBS followed by centrifugation at 2000 rpm for 2 min. Cell pellets were then resuspended in 200 uL PBS and transferred to a 96 well U-bottom plate for staining. Cells were then incubated with Zombie Aqua fixable viability stain (1:1000, Biolegend) for 15 min prior to staining with primary antibodies diluted in PBS+2% Heat Inactivated FBS (FACS buffer) for 30 min on ice (see Antibody table). Cells were then washed twice with FACS buffer prior to analysis on a LSR II analytical flow cytometer.

For FACS analysis of tumor bearing lung tissue, KP or KP; K^b^Strep mice (n=3 each) were euthanized 12 weeks post tumor initiation. 2 minutes prior to sacrifice, intravascular staining antibody (anti-CD45-APC-eFluor786, Biolegend) was injected retro-orbitally to stain circulating immune cells. Mice were then euthanized with cervical dislocation, and the lungs were removed and placed on ice. 100-200 mg of tumor bearing lung tissue was then thoroughly minced with Noyes scissors, prior to incubation with digestion buffer (HBSS supplemented with 5% HI FBS, 125 U/mL collagenase IV (Worthington) and DNase (Roche) with rotation for 30 min at 37 °C. Cell suspensions were then macerated through 70 µM cell strainers (Corning) and centrifuged at 500 × g for 5 min. Cell pellets were resuspended in 1 mL ACK Lysis Buffer (Gibco) and incubated at room temperature for 5 min before quenching with 10 mL RPMI + 10% HI FBS and centrifugation at 500 × g for 5 min. Cell pellets were then resuspended in 200 uL FACS buffer and transferred to 96 well U bottom plates for staining with Fc block (BD Biosciences), zombie fixable viability stain for 20 min, and primary antibody cocktail for 30 min on ice. Cells were then washed three times with FACS buffer before fixation overnight with FoxP3/Transcription Factor Staining Buffer Set (eBioscience) according to manufacturer’s instructions prior analysis on a Fortessa flow cytometer. All flow cytometry data was analyzed in FlowJo v10.

### Affinity Purification of H2-K^b^ with Streptactin

Whenever possible, great care was taken to keep samples ice cold at all times to maintain MHC-I complex stability. For cultured cells, ∼6 × 10^7^ cells were used for each replicate (4 × 15 cm dishes). In the culture dish, cells were washed twice with PBS, prior to lysis with 2 mL MHC Extraction Buffer (MEB: 20 mM Tris pH 8.0, 100 mM NaCl, 1 mM EDTA, 1% Triton-X100, 60 mM octyl-glucopyranoside, 6 mM MgCl_2_, and 1X HALT protease inhibitors (Pierce). Cell lysates were then transferred to 2 mL microcentrifuge tubes and supplemented with 20 U benzonase and 10 U avidin (to block endogenously biotinylated protein) prior to incubating with rotation at 4 °C. Lysates were then cleared by centrifugation at 16,000 × g for 15 min prior to incubation with MagStrep Type 3 Streptactin XT beads (IBA Biosciences).

For pancreatic and lung tissues, tumor bearing tissue was dissected and immediately lysed or flash frozen in liquid N_2_ for later processing. For lysis, fresh or frozen tissue was quickly minced with Noyes scissors and transferred to a 7 mL glass Dounce homogenizer (Sigma), precooled on ice. 4 mL MEB was then added, and the tissue was thoroughly homogenized with 10-20 passes of a loose-fitting pestle followed by 5-10 passes of a tight-fitting pestle. Tissue homogenates were transferred to 5 mL centrifuge tubes (Eppendorf) and supplemented with 20 U benzonase and 10 U avidin prior to incubating with rotation at 4 °C for 20 min and subsequent removal of debris by centrifugation at 16,000 × g for 15 min.

Prior to incubating with cleared lysate, Streptactin beads were equilibrated by magnetizing and washing 1x with MEB. For each cell culture sample or tissue sample, 1 mL of bead suspension (50 uL bed volume) was used. Equilibrated beads were then added to cleared lysates and incubated with rotation at 4 °C for 1-3 hrs. After incubation, beads were washed 2 × with MEB, 2 × with TBS, and 2x with 20 mM Tris. On the last wash, suspended beads were transferred to a new Lo-Bind microcentrifuge tube prior to elution. Strep tagged H2-K^b^ was then eluted from the Streptactin resin by adding 400 uL of 0.5X Buffer BXT (IBA Biosciences) and incubating on ice for 20-30 min with occasional flicking of the tube to maintain the beads in suspension. Beads were then magnetized, and the supernatant was transferred to a new Lo-Bind microcentrifuge tube. Biotin, H2-K^b^ heavy chain, and B2m light chain were then precipitated by adding 1% TFA slowly while gently vortexing the elution. The fluffy white precipitate was then pelleted by centrifugation at 20,000 × g for 10 min. Supernatants containing liberated MHC-I peptides were then directly aspirated into 8 ug binding capacity C18 solid phase extraction pipette tips. Tips were preequilibrated with 50% Acetonitrile (ACN) and 0.1 % formic acid according to manufacturer’s instructions. The 400 uL of eluted material was loaded, 20 uL at a time until the entire volume of the elution was passed over the C18 sorbent. Tips were then washed twice with 5% ACN/0.1% FA prior to elution in 10 uL of 30% ACN/0.1% FA. Desalted peptides were then lyophilized prior to LC-MS/MS analysis.

### Antibody Immunoprecipitation of MHC-I

Peptide MHC isolation was performed as described previously^33^. Healthy lung tissue or tumor bearing lung tissue was homogenized and cleared as described for Streptactin purification. Per sample, 1 mg of Anti-H2-K^b^ (clone Y3, BioXCell) was bound to 20 μL (bed volume) FastFlow Protein A Sepharose beads (GE Healthcare) by incubating for 1 hour at 4°C. Beads were then washed with lysis buffer samples were incubated for 4 hours rotating at 4°C. Beads were then centrifuged at 2,000 rpm, washed twice with 1× TBS, and eluted with 10% acetic acid at room temperature. Eluate was then filtered using 10 kDa MWCO spin filters (PALL Life Science), which were passivated with 0.1% BSA and acidified with 10% acetic acid prior to filtration. Filtered peptides were then further purified with 8 µg binding capacity C18 tips (Pierce) prior to LC/MS-MS analysis.

### Liquid Chromatography Tandem Mass Spectrometry

pMHC samples were analyzed using an Exploris 480 Hybrid Quadrupole-Orbitrap mass spectrometer (Thermo Scientific) coupled with an UltiMate 3000 RSLC Nano LC system (Dionex), Nanospray Flex ion source (Thermo Scientific), and column oven heater (Sonation). Samples were resuspended in 0.1% formic acid and directly loaded onto an analytical capillary chromatography column with an integrated electrospray tip (∼1 μm orifice), prepared and packed in house (50 μm ID × 15 cm & 1.9 μM C18 beads, ReproSil-Pur). Fifty percent of the pMHC elutions was analyzed in a given LC-MS/MS analysis.

Peptides were eluted using a linear gradient with 6-25% buffer B (70% Acetonitrile, 0.1% formic acid) for 53 minutes, 25-45% for 12 minutes, 45-97% for 3 minutes, hold for 1 minute, and 97% to 3% for 1 minute.

Standard mass spectrometry parameters were as follows: spray voltage, 2.0-2.5 kV; no sheath or auxiliary gas flow; heated capillary temperature, 275 °C. The Exploris was operated in data dependent acquisition (DDA) mode. Full scan mass spectra (350-1200 m/z, 60,000 resolution) were detected in the orbitrap analyzer after accumulation of 3E6 ions (normalized AGC target of 300%), automatic maximum injection time (IT). For every full scan, MS^2^ spectrum were collected during a 3 second cycle time. Ions were isolated (0.4 m/z isolation width) for a maximum IT of 150/250 ms or 1E5/7.5E4 ions (100%/75% normalized AGC target) and fragmented by HCD with 30% collision energy at a resolution of 60,000. Charge states < 2 and > 4 were excluded, and precursors were excluded from selection for 30 seconds if fragmented n=2 times within 20 second window.

### Mass Spectrometry data analysis

All mass spectra were analyzed with Proteome Discoverer (PD, version 2.5) and searched using Mascot (version 2.4) against the mouse SwissProt database (2021_02) supplemented with a list of murine ORFs previously identified by ribosome profiling (www.sorfs.org) Peptides were searched with no enzyme, variable OxM, Peptides were further filtered according to the following criteria: ion score ≥ 15, search engine rank = 1. Results from technical replicates of each sample analysis were combined. Area under the curve (AUC) quantitation was performed using the minora feature detector in PD with match between runs enabled and filtered for ion score ≥ 15, search engine rank = 1. Abundances were averages across technical and biological replicates.

### Immunofluorescence of Lung and Pancreas Tissue

Tumor bearing lung or pancreas tissue was manually dissected and embedded in optimal cutting temperature (OCT) compound and slowly frozen on dry ice. Frozen tissue sections were stored at -80 °C until sectioning. On the day of sectioning, frozen tissue was allowed to equilibrate to -20 °C in the cryostat for at least 1 hour. 8 µM sections were then cut and transferred to microscope slides (Fisher) prior to fixation in 100% acetone at -20 °C for 10 min and dried overnight at room temperature. Slides were then stored at -20 °C until staining.

Tissue sections were rehydrated with PBS for 5-10 mins and then blocked in PBS supplemented with 5% BSA for 45 minutes. Primary antibodies were then added at the indicated dilutions (see Antibody Table) for 1 hour at 25 °C. Slides were then washed 4 times with PBS + BSA, and incubated with fluorescently labeled secondary antibodies where indicated for 1 hour at 25 °C. Stained sections were then washed 3 times with PBS, incubated with DAPI for 5 min, washed once with PBS, and then mounted with Prolong Diamond AntiFade Mountant (Thermo). Slides were scanned on a VERSA 8 slide scanner (Leica) prior to analysis in ImageScope v64 and ImageJ.

### Immunoblotting

For organoids, cells were dissociated with TrypLE, washed once with PBS, and then lysed in cell lysis buffer (CLB, 50 mM Tris pH 7.5, 150 mM NaCl, 1 mM EDTA, 1% Triton-X100, 0.1% SDS, 1X HALT protease inhibitors). For monolayer cultures, media was aspirated cells were washed twice with PBS prior to lysis with CLB. Protein concentration was quantified with BCA (Pierce) and samples were loaded onto 4-12% Bis-Tris SDS-PAGE gels (Invitrogen) and electrophoresed at 150V until the loading dye reached the bottom of the gel. Gels were then transferred onto 0.45 uM nitrocellulose membranes overnight at 20V in a cold room. Blots were blocked with PBST (PBS + 0.5% Tween20) + 5% milk for 30 min at room temperature, incubated with primary antibody for 1 hour at room temperature, washed 4 times in PBST, incubated with HRP-conjugated secondary antibody for 1 hour at room temperature, and then washed 4 times in PBST. Blots were developed with Clarity or Clarity-Max ECL substrated (BioRad) and imaged on a ChemiDoc Gel Imaging System (BioRad).

### RT-PCR

RNA was isolated from organoid cultures by aspirating media, and then resuspending the Matrigel domes in 1 mL Trizol (Ambion). RNA was then extracted from the homogenate using the PureLink RNA Mini Kit (Ambion) according to manufacturer’s instructions with with the final elution step in 40 µL of nuclease free water. Isolated RNA was then quantified on a NanoDrop 2000 (Thermo). 1 µg of total RNA was then added to a cDNA synthesis reaction using a High Capacity cDNA Reverse Transcription Kit (Applied Biosciences) according to manufacturer’s instructions with random hexamer’s as primers and including RNase inhibitor in the reaction. cDNA was then diluted 1:10 with nuclease free water prior to adding 1 µL to a PCR reaction containing 500 uM forward and reverse primer and 1X Q5 Master Mix (NEB). Reactions were cycled 35 times with annealing temperatures calculated by NEB Tm calculator prior to loading on 2% agarose gels. DNA bands were imaged by staining with ethidium bromide and UV transillumination.

### Ribosome Protected Fragment Isolation with RiboLace

Ribosome profiling was performed using a previously described method using Puromycin conjugated magnetic beads to isolated actively translating ribosomes (Clamer et. al. 2018 Cell Reports). To obtain cellular material for Ribosome sequencing, organoid cultures of normal alveolar type 2 cells or organoids derived from 12 week KP tumors were transiently adapted to monolayer culture by washing tissue cultures dishes with a 10% Matrigel solution prior to plating cells in either complete lung organoid medium for AT2 cells or lung organoid base medium for tumor organoids. After 48 hours, cells were treated with 100 µM cycloheximide for 5 min and immediately transfer to ice. Plates were then washed once with 10 mL PBS + 100 µM cycloheximide followed by direct lysis and thorough mechanical disruption in the plate according to the RiboLace Protocol (Immagina Biotech). Lysates were then cleared with centrifugation at 20,000 × g for 15 min. Nucleic acid content was quantified with a NanoDrop and each lysate was normalized to 0.3 a.u._260nm_ in 150 µL of W-buffer. 0.3 µL of SS solution and diluted RNAse Nux solution (5 µL of 1:66.7 dilution). RNase digestion was then stopped with 0.5 µL SUPERaseIn for 10 min on ice.

RiboLace beads were prepared according to RiboLace kit instructions. Briefly, per sample, 90 µL of beads were magnetized, washed once with buffer OH, once with nuclease free water, twice with B-buffer, and then resuspended in 30 µL RiboLace probe followed by incubation shaking for 1 hour at room temperature. Beads were then passivated with 3 µL PEG for 15 min at room temperature and washed twice with 500 µL of buffer W.

Prepared beads were then resuspended in RNAse digested lysates, and incubated with slow rotation for 70 min at 4 °C. Beads were then washed twice with 500 µL of Buffer-W before protease digestion by addition of 20 µL of SDS and 5 µL of proteinase K and incubating for 75 min at 37 °C. RNA was then extracted with Phenol-Chloroform, precipitated overnight with isopropanol, and then electrophoresed on denaturing 15% TBE-Urea gels. Gel pieces were excised from the region according for 20-40 nucleotides to enrich for ribosome protected fragments (RPFs). Gel pieces were then crushed by centrifugation through a punctured 0.5 mL tube placed in a 1.5 mL microcentrifuge tube. Gel debris was then incubated overnight rotating in 400 µL of gel elution buffer (20 mM Tris pH 7.5, 250 mM sodium acetate, 1 mM EDTA, 0.25% SDS). RNA was precipitated from the elutions by adding 700 µL of isopropanol and 1.5 µL of GlycoBlue (Thermo) followed by overnight incubation at -80°C. RNA was then pelleted by centrifugation at 20,000 × g for 30 min at 4 °C. The RNA pellet was then washed with 70% EtOH, air dried for 2-3 min and resuspended in 10 µL of nuclease free water.

Dephosphorylation of RPFs was performed by adding 23 µL nuclease free water, 5 µL T4 PNK buffer (NEB), 1 µL SUPER In (Invitrogen), and 1 µL T4 PNK (NEB) followed by incubation for 60 min at 37 °C, then 10 min at 70 °C, and cooled to room temperature. RNA was then precipitated by adding 39 µL nuclease free water, 10 µL 3M sodium acetate, 150 µL isopropanol, and 1 µL GlycoBlue and incubated at -80 °C overnight. RNA was pelleted by centrifugation at 20,000 × g for 30 min, washed with 70% EtOH and then resuspended in 7 µL of nuclease free water.

### RPF Sequencing Library Preparation

Illumina sequencing libraries of isolated RPFs was performed with the SMARTer smRNA-Seq Kit for Illumina (Takara Bio). Library preparation was performed according to the manufacturer’s protocol with the following parameters. Polyadenylation was performed with 7 µL of the resuspended RPFs, with addition of 0.25 µL of Poly(A) Polymerase (2 U/µL), 0.25 µL RNase inhibitor (40 U/µL) and 2.5 µL of smRNA Mix 1 with no supplemental ATP. Reactions were incubated at 16 °C for 5 min and then immediately placed back on ice. cDNA synthesis was performed by adding 1 µL of smRNA dT Primer, incubating for 3 min at 72 °C and then transferring to ice. Next, 6.5 µL of smRNA Mix 2, 0.5 µL RNAse inhibitor, and 2 µL PrimeScript RT was added followed by incubation for 60 min at 42 °C, then 10 min at 70 °C, and transferred to ice. cDNA was amplified by adding 24 µL nuclease free water, 50 µL 2X SeqAmp PCR Buffer, 2 µL SeqAmp DNA Polymerase, and 2 µL each of indexed forward and reverse primer. Reactions were cycled 11 times at 98 °C for 10 s, 60 °C for 5 s, and 68 °C for 10 s. Amplified libraries were purified using a NucleoSpin Gel and PCR Clean-Up Kit (Macherey-Nagel) and size selected by electrophoresis on non-denaturing 6% TBE-Urea gels. Bands according to ∼175 bp were excised and crushed, and eluted with 400 µL of GEB. Eluates were then concentrated in a SpeedVac to ∼100 µL and purified with the DNA Clean and Concentrator Kit (Zymo). Libraries were sequenced on a HiSeq 2000 at the BioMicroCenter Core Facility at MIT with 50 nt read length.

### RiboSeq Data Analysis

Raw fastq files from each RiboSeq library were trimmed according to recommendations in the SMARTer smRNA-Seq protocol. Fastq files from each sample type were then merged prior to submission to the RiboToolKit server (http://rnabioinfor.tch.harvard.edu/RiboToolkit/) using a 26-38 nt size filter and a predicted ORF cutoff of p < 0.05. Data were then downloaded and visualized in R. For RPKM values presented in Figure 3 and Figure 6, total RPKM across the entire transcript was used.

### Hsp90 Inhibitor Treatment

Hsp90 inhibitor (NVP-HSP990, Selleck Chemical) was administered in the drinking water. The average water consumption of the mice was calculated by measuring water bottle weight before and after 72 hours of housing to determine the average consumption per mouse, per day. Across all experiments, C57Cl/6 mice consumed approximately 4 mL every day. Using these water consumption values, and mouse weight, a 4 mg/mL stock solution of NVP-HSP990 (in 100% PEG400) was diluted directly into the drinking water to achieve a target dose of 0.5 mg/kg/day. Hsp90i treatment began 8 weeks post tumor induction and water was replaced twice per week for 4 weeks prior to sacrifice.

### Data Analysis, Statistics, and Visualization

All quantitative data was processed and visualized in R (R version 4.0.2, RStudio version 1.3.959). Peptides from PDAC or LUAD samples were concatenated and organized in Microsoft Excel with predicted affinity values calculated by NetMHCPan 4.1. Following import into R, peptide lists were filtered for length (8-11 aa) and predicted affinity (<1000 nM) before further analysis. Non-metric multidimensional scaling (nmds) analysis was performed using the published R script from Sarkizova et. al. and applied to 8- and 9-mer peptides separately. Peptide clusters were empirically chosen through hierarchical clustering of the calculated nmds matrix. Peptide motifs were generated with the ggseqlogo package in R. All boxplots are in the Tukey style with the horizontal mark at the median, the box extending from quartile 1 to quartile 3, and whiskers extending 1.5 x the interquartile range from the box.

Bulk RNA sequencing was adapted from Chuang et. al. by taking the mean expression of each gene across all normal, early, nonmet, and metastatic primary tumor samples prior to cross comparison to peptide data.

Single cell RNA sequencing data (scRNAseq) was analyzed with Seurat (version 4.0.1). Cells by genes matrices were downloaded from the Tabula Muris (healthy lung and pancreas) or from the Broad single cell RNA seq Portal (KP scRNAseq data). Data were filtered with a gene count cutoff of 1100 prior to normalization, scaling, and dimensionality reduction with UMAP according to standard Seurat pipelines (https://satijalab.org/seurat/). Custom modules were added with the AddModuleScore function according to gene lists from peptide data or published reports. Volcano plots for comparing signatures across cell types in the healthy lung data were calculated with two-tailed Student’s t-test for all pairwise comparisons of Normal/Ab/Strep signatures within each cell type with a Bonferroni multiple comparisons adjustment. The median score for each signature was then used to calculate the Log_2_ fold change and *P* values were transformed to -Log_10_(*P*) for plotting. Pearson correlations between gene signatures were calculated by AddModuleScore in Seurat, extraction of metadata, conversion of the data to a matrix containing cells x signatures, and calculation of Pearson correlation for signatures across cells. For correlation of peptide signatures to all genes, the scaled data from Seurat was extracted to a matrix, and the Late/Tumor-Unique Peptide signature was appended onto the matrix. Pearson correlation was then calculated between the peptide signature versus each individual gene. Genes were then ranked by correlation and analyzed with preranked gene set enrichment in GSEA (Broad Institute).

Gene Ontology analysis was performed on Gene Lists in StringDb (www.stringdb.org) with high confidence interaction cutoffs. Gene Set enrichment analysis (Figure 2) was performed with the GSEA software package using a preranked gene list according to correlation to the Late or Tumor-Unique peptide signature.

Subcellular compartment analysis was performed by extracting compartment locations from UniProt for source proteins which had available localization data. Comparison of subcellular distributions was performed with Fisher’s Exact test with Monto Carlo simulation. Protein Length information was also extracted from Uniprot for each source protein. Thermal stability and protein half-life were obtained from published data sets^30^.

Statistical analyses were performed in R. To compare median fluorescence intensity from flow cytometry data, two-tailed Student’s t-tests were performed for all pairwise comparisons. For comparisons of gene expression or signature scores an unpaired two sample Wilcoxon Test (Mann-Whitney *U*) was performed. For analysis of peptide abundance distributions in Veh and Hsp90i treated samples, Kolmogorov-Smirnov Tests were used.

## Data Availability

All mass spectrometry data have been deposited to the ProteomeXchange Consortium via the PRIDE partner repository with the dataset identifier PXD026337. Raw ribosome sequencing data has been submitted to the Gene Expression Omnibus with dataset identifier [GEOXXXXX].

## References

1. Schumacher, T. N. & Schreiber, R. D. Neoantigens in cancer immunotherapy. Science 348, 69–74 (2015).

2. Jhunjhunwala, S., Hammer, C. & Delamarre, L. Antigen presentation in cancer: insights into tumour immunogenicity and immune evasion. Nat Rev Cancer 1–15 (2021) doi:10.1038/s41568-021-00339-z.

3. Keenan, T. E., Burke, K. P. & Allen, E. M. V. Genomic correlates of response to immune checkpoint blockade. Nat Med 25, 389–402 (2019).

4. Dersh, D., Hollý, J. & Yewdell, J. W. A few good peptides: MHC class I-based cancer immunosurveillance and immunoevasion. Nat Rev Immunol 1–13 (2020) doi:10.1038/s41577-020-0390-6.

5. Ghorani, E. et al. The T cell differentiation landscape is shaped by tumour mutations in lung cancer. Nat Cancer 1, 546–561 (2020).

6. Abelin, J. G. et al. Mass Spectrometry Profiling of HLA-Associated Peptidomes in Mono-allelic *Cell*s Enables More Accurate Epitope Prediction. Immunity 46, 315–326 (2017).

7. Wells, D. K. et al. Key Parameters of Tumor Epitope Immunogenicity Revealed Through a Consortium Approach Improve Neoantigen Prediction. Cell 183, 818–834.e13 (2020).

8. Chong, C. et al. Integrated proteogenomic deep sequencing and analytics accurately identify non-canonical peptides in tumor immunopeptidomes. Nat Commun 11, 1293 (2020).

9. Yadav, M. et al. Predicting immunogenic tumour mutations by combining mass spectrometry and exome sequencing. Nature 515, 572–576 (2014).

10. Granados, D. P. et al. Impact of genomic polymorphisms on the repertoire of human MHC class I-associated peptides. Nat Commun 5, 3600 (2014).

11. Sarkizova, S. et al. A large peptidome dataset improves HLA class I epitope prediction across most of the human population. Nat Biotechnol 38, 199–209 (2019).

12. Stopfer, L. E., Mesfin, J. M., Joughin, B. A., Lauffenburger, D. A. & White, F. M. Multiplexed relative and absolute quantitative immunopeptidomics reveals MHC I repertoire alterations induced by CDK4/6 inhibition. Nat Commun 11, 2760 (2020).

13. Khodadoust, M. S. et al. Antigen presentation profiling reveals recognition of lymphoma immunoglobulin neoantigens. Nature 543, 723–727 (2017).

14. Hu, Z. et al. Personal neoantigen vaccines induce persistent memory T cell responses and epitope spreading in patients with melanoma. Nat Med 27, 515–525 (2021).

15. DuPage, M., Dooley, A. L. & Jacks, T. Conditional mouse lung cancer models using adenoviral or lentiviral delivery of Cre recombinase. Nat Protoc 4, 1064–1072 (2009).

16. Jamal-Hanjani, M. et al. Tracking the Evolution of Non–Small-Cell Lung Cancer. New Engl J Medicine 376, 2109–2121 (2017).

17. Roe, J.-S. et al. Enhancer Reprogramming Promotes Pancreatic Cancer Metastasis. Cell 170, 875–888.e20 (2017).

18. Chiou, S.-H. et al. Pancreatic cancer modeling using retrograde viral vector delivery and in vivo CRISPR/Cas9-mediated somatic genome editing. Gene Dev 29, 1576–1585 (2015).

19. Schaum, N. et al. Single-cell transcriptomics of 20 mouse organs creates a Tabula Muris. Nature 562, 367–372 (2018).

20. Stegemann-Koniszewski, S. et al. Alveolar Type II Epithelial Cells Contribute to the Anti-Influenza A Virus Response in the Lung by Integrating Pathogen- and Microenvironment-Derived Signals. Mbio 7, e00276–16 (2016).

21. Schuster, H. et al. A tissue-based draft map of the murine MHC class I immunopeptidome. Sci Data 5, 180157 (2018).

22. Marjanovic, N. D. et al. Emergence of a High-Plasticity Cell State during Lung Cancer Evolution. Cancer Cell 38, 229–246.e13 (2020).

23. LaFave, L. M. et al. Epigenomic State Transitions Characterize Tumor Progression in Mouse Lung Adenocarcinoma. Cancer Cell 38, 212–228.e13 (2020).

24. Dongre, A. & Weinberg, R. A. New insights into the mechanisms of epithelial–mesenchymal transition and implications for cancer. Nat Rev Mol Cell Bio 20, 69–84 (2019).

25. Yewdell, J. W., Dersh, D. & Fåhraeus, R. Peptide Channeling: The Key to MHC Class I Immunosurveillance? Trends Cell Biol 29, 929–939 (2019).

26. Weinzierl, A. O. et al. Distorted Relation between mRNA Copy Number and Corresponding Major Histocompatibility Complex Ligand Density on the Cell Surface* S. Mol Cell Proteomics 6, 102–113 (2007).

27. Chuang, C.-H. et al. Molecular definition of a metastatic lung cancer state reveals a targetable CD109–Janus kinase–Stat axis. Nat Med 23, 291–300 (2017).

28. Clamer, M. et al. Active Ribosome Profiling with RiboLace. Cell Reports 25, 1097–1108.e5 (2018).

29. Chen, J. et al. Pervasive functional translation of noncanonical human open reading frames. Science 367, 1140–1146 (2020).

30. Perrin, J. et al. Identifying drug targets in tissues and whole blood with thermal-shift profiling. Nat Biotechnol 38, 303–308 (2020).

31. Bassani-Sternberg, M., Pletscher-Frankild, S., Jensen, L. J. & Mann, M. Mass Spectrometry of Human Leukocyte Antigen Class I Peptidomes Reveals Strong Effects of Protein Abundance and Turnover on Antigen Presentation. Mol Cell Proteomics 14, 658–673 (2015).

32. Brodsky, J. L. Cleaning Up: ER-Associated Degradation to the Rescue. Cell 151, 1163–1167 (2012).

33. Jaeger, A. M. et al. Rebalancing Protein Homeostasis Enhances Tumor Antigen Presentation. Clin Cancer Res 25, 6392–6405 (2019).

34. Taipale, M., Jarosz, D. F. & Lindquist, S. HSP90 at the hub of protein homeostasis: emerging mechanistic insights. Nat Rev Mol Cell Bio 11, 515–528 (2010).

35. Savitski, M. M. et al. Multiplexed Proteome Dynamics Profiling Reveals Mechanisms Controlling Protein Homeostasis. Cell 173, 260–274.e25 (2018).

36. Karras, G. I. et al. HSP90 Shapes the Consequences of Human Genetic Variation. Cell 168, 856–866.e12 (2017).

37. Jarosz, D. F., Taipale, M. & Lindquist, S. Protein Homeostasis and the Phenotypic Manifestation of Genetic Diversity: Principles and Mechanisms. Annu Rev Genet 44, 189–216 (2010).

38. Molina, D. M. et al. Monitoring Drug Target Engagement in Cells and Tissues Using the Cellular Thermal Shift Assay. Science 341, 84–87 (2013).

39. Stevanović, S. et al. Landscape of immunogenic tumor antigens in successful immunotherapy of virally induced epithelial cancer. Science 356, 200–205 (2017).

40. Sahin, U. et al. An RNA vaccine drives immunity in checkpoint-inhibitor-treated melanoma. Nature 585, 107–112 (2020).

41. Zitvogel, L., Perreault, C., Finn, O. J. & Kroemer, G. Beneficial autoimmunity improves cancer prognosis. Nat Rev Clin Oncol 1–12 (2021) doi:10.1038/s41571-021-00508-x.

42. Nelson, C. E. et al. Robust Iterative Stimulation with Self-Antigens Overcomes CD8+ T Cell Tolerance to Self- and Tumor Antigens. Cell Reports 28, 3092–3104.e5 (2019).

43. Vonderheide, R. H. CD40 Agonist Antibodies in Cancer Immunotherapy. Annu Rev Med 71, 1–12 (2019).

44. Cuevas, M. V. R. et al. Most non-canonical proteins uniquely populate the proteome or immunopeptidome. Cell Reports 34, 108815 (2021).

45. Erhard, F. et al. Improved Ribo-seq enables identification of cryptic translation events. Nat Methods 15, 363–366 (2018).

46. Ouspenskaia, T. et al. Thousands of novel unannotated proteins expand the MHC I immunopeptidome in cancer. Biorxiv 2020.02.12.945840 (2020) doi:10.1101/2020.02.12.945840.

47. Raposo, B. et al. T cells specific for post-translational modifications escape intrathymic tolerance induction. Nat Commun 9, 353 (2018).

48. Griffin, G. K. et al. Epigenetic silencing by SETDB1 suppresses tumour intrinsic immunogenicity. Nature 1–6 (2021) doi:10.1038/s41586-021-03520-4.

49. Kalaora, S. et al. Identification of bacteria-derived HLA-bound peptides in melanoma. Nature 592, 138–143 (2021).

